# A scalable transposon mutagenesis system for non-model bacteria

**DOI:** 10.64898/2025.12.22.696024

**Authors:** Charlie Gilbert, Julia Leung, Alexander Crits-Christoph, Shinyoung Clair Kang, Ariela Esmurria, Kathrin Fenn, Stephanie L. Brumwell, Zaira Martin-Moldes, Kerrin Mendler, Tyler P. Barnum, Mary-Anne Nguyen, Henry H. Lee, Nili Ostrov

**Affiliations:** Cultivarium, 490 Arsenal Way, Watertown MA 02472, USA

## Abstract

Transposon mutagenesis enables genome-wide interrogation of gene function with a single self-contained genetic construct. However, its application to non-model bacteria remains limited because transposition efficiency depends on multiple host factors that are difficult to predict *a priori* including transposase activity, antibiotic resistance marker performance, and regulatory element compatibility. Here, we present a scalable system to identify functional transposon configurations in non-model bacteria through pooled library screening. We selected 18 promoters across multiple bacterial phyla to independently drive expression of the transposase and antibiotic resistance marker, generating 324 promoter combinatorial variants for each of six antibiotic resistance markers. We developed a high-throughput, automated workflow to deliver all 1,944 mariner-based transposon variants in a single experiment and applied this to 92 non-model bacteria spanning multiple phyla. From this, we identified functional transposons for 43 strains, with high-level mutagenesis (10^2^-10^4^ unique insertions) in 13 species, including seven with no previously described transposon mutagenesis. We then expanded to a dual-transposase system, mariner or Tn5, and devised a single transposon insertion sequencing method for high-throughput screening of 3,888 configurations. To demonstrate the practical utility of our screening approach, we used a top-performing variant to generate a genome-wide transposon mutant library for *Comamonas testosteroni* KF-1, a bacterium that metabolizes plastic- and lignin-derived polymers. We assayed this *C. testosteroni* mutant library to identify enzymatic pathways, transporter genes, and regulators essential for the metabolism of plastics-associated monomer terephthalate and lignin-associated monomer 4-hydroxybenzoate. Together, this work establishes a scalable approach to construct and identify genetic perturbation systems in non-model bacteria, expanding our ability to systematically probe gene function across the bacterial tree of life.

## Introduction

The investigation of gene function remains laborious and slow in most microbial systems, particularly those that have little supporting literature. Transposon mutagenesis has emerged as a mainstay approach for genome-scale interrogation of gene function: a single genetic construct generates hundreds of thousands of mutants, which can be assayed across growth conditions in a single pooled experiment^1–4^.

Beyond mutagenesis, transposons offer a route to establishing genetic tractability in non-model microbes^5–8^. Alternative approaches exist, such as screening for compatible origins of replication^9^, but unknown host factors make it difficult to generalize across organisms. Transposons, by contrast, are defined by a small number of genetic parts: a transposase and a selectable marker, each requiring functional expression in the host. The factors governing this expression are constrained predominately by promoters, ribosome binding sites (RBS), and codon usage – making transposons simpler to optimize.

However, establishing genome-wide transposon mutagenesis in a non-model microbe remains challenging. An optimal transposon requires balanced expression of both the transposase and selectable marker, as well as low insertional bias to ensure even genomic coverage. These parameters depend on multiple factors including transposase type, effective transposon expression, and selection marker efficiency^10–13^.

We and others have demonstrated that multiplexed sequencing assays can identify functional genetic parts from a large combinatorial parameter spaces^9,14–21^. A previous study dubbed “magic pools” reported screening of four transposon variant libraries in six bacterial strains, using the best performing library members for genome-wide transposon mutagenesis^11^. Building upon this, we generated transposon variant libraries with broader antibiotic coverage (six versus two resistance marker types), standardized promoter combinations between libraries, and developed a high-throughput, automated assay for rapid identification of functional transposons. We applied this system to 92 non-model bacteria to identify functional transposons for 43 strains, including high-confidence transposon constructs for 13 species (10^2^-10^4^ unique insertions). Furthermore, we utilized this system to generate a genome-wide mutant library for *Comamonas testosteroni* KF-1, a bacterium capable of degrading plastic- and lignin-derived compounds. We assayed this library to identify enzymatic pathways, transporters, and regulators required for growth on terephthalate (TER) and 4-hydroxybenzoate (4HB).

## Results and Discussion

### Design and validation of transposon variant libraries

To enable the identification of plasmid constructs for efficient transposon mutagenesis in non-model bacteria, we built libraries of transposon variants specifically designed for pooled screening (**Figure 1A**). Each transposon construct contains a transposase gene and an antibiotic resistance (ABR) marker. The libraries, which we refer to as ‘transposon variant libraries’ are composed of variable transposase and ABR genes as well as variable promoters for both. Transposon variant libraries are designed to be delivered in pooled fashion to target bacteria with subsequent antibiotic selection and sequencing-based identification of the optimal combination of parts for high transposition efficiency in a process we refer to as ‘transposon variant screening’.

**Figure 1.**
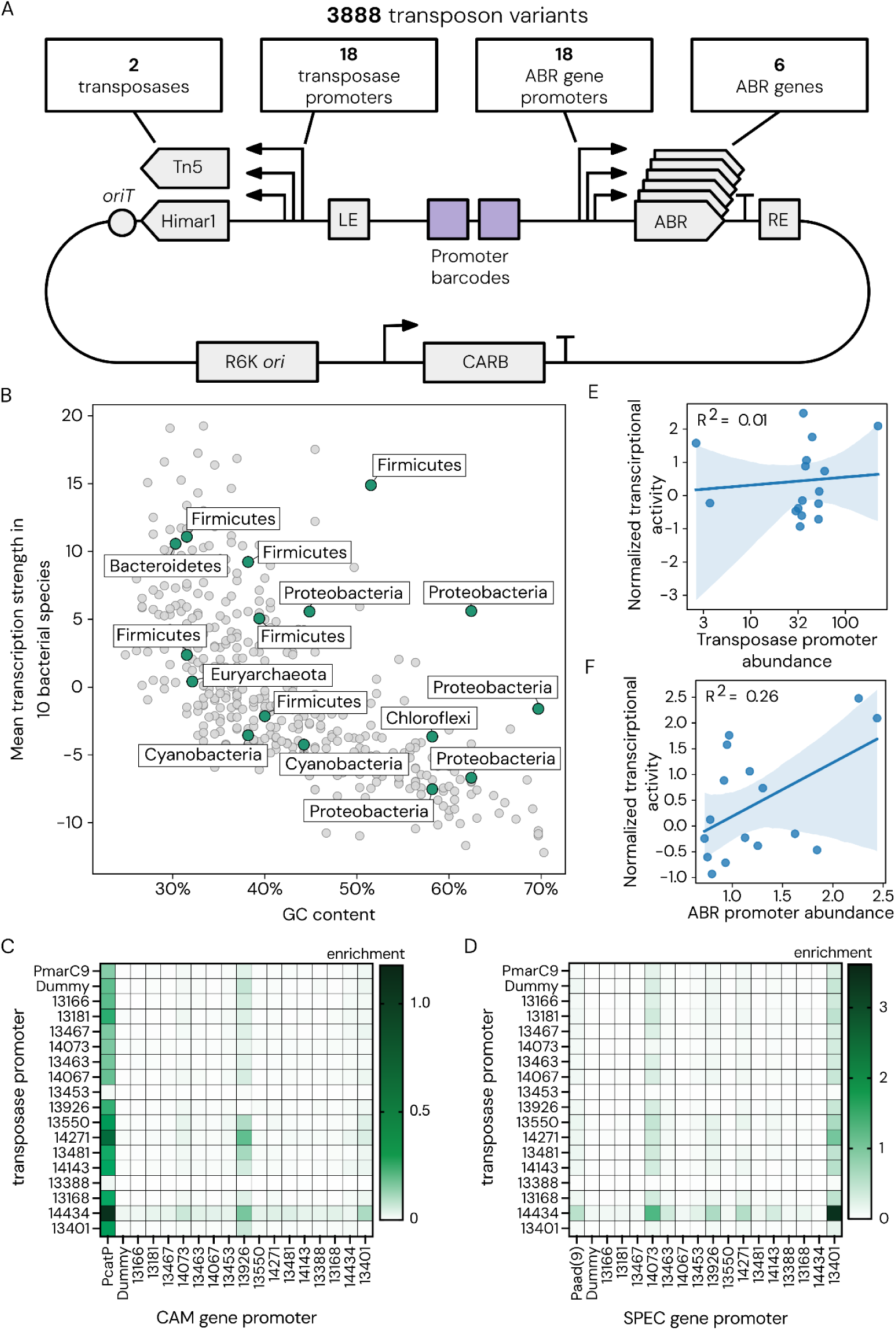
Design and validation of the transposon variant libraries. (**A**) Schematic of the transposon variant library components. LE, left mosaic end; RE, right mosaic end; *oriT*, origin of transfer; ABR, antibiotic resistance. (**B**) Mean transcription strength and GC content (%) of 16 promoters selected for inclusion in the transposon variant library (green). Transcription strength for 421 promoter sequences was measured across 10 bacterial strains by Yim et al. 2019^25^. (**C**) Himar1-only library variant performance in *E. coli* DH10B, with either the *catP* (CAM) or (**D**) *aadA(9)* (SPEC) ABR gene. Relative transposition efficiency (enrichment) was measured by Tn-seq after pooled conjugative delivery. (**E**) Correlation between previously published^25^ transcriptional strength of promoters and their transposition efficiencies in *E. coli* when driving the Himar1 transposase. (**F**) Correlation between previously published^25^ transcriptional strength of promoters and their transposition efficiencies in *E. coli* when driving the *catP* ABR.

We selected two of the most highly used transposases for bacterial mutagenesis, Himar1^22^ (a hyperactive variant of a mariner transposase) and Tn5^23,24^, and six commonly used ABR markers (**Supplementary Table 1**): chloramphenicol (CAM), spectinomycin (SPEC), kanamycin (KAN), gentamicin (GEN), tetracycline (TET), and erythromycin (ERM)^20^. Finally, we selected a functionally-diverse set of 16 promoter-5’UTR (‘promoter’) sequences to drive expression of both the transposase and ABR marker gene.

Promoter sequences were selected from a range of source organisms, GC content, and activity profiles. We constrained the total number of promoter variants tested to 16, as increased library sizes result in dilution of individual functional components within the library, requiring higher transformation efficiencies for detection. In an effort to include promoters with broad host ranges that are less dependent on specific host factors, we selected from a dataset of 421 promoters associated with mobile genetic elements, for which transcription strengths were previously characterized in cell-free systems of 10 bacterial strains^25^. We developed an algorithm to prioritize 16 promoters depleted in potential restriction enzyme cut sites, with a range of GC contents, and originating from six phyla: Firmicute (Bacillota), Bacteroidetes (Bacteroidota), Chloroflexi (Chloroflexota), Cyanobacteria (Cyanobacteriota), Euryarchaeota (Halobacteriota), and Proteobacteria (Pseudomonadota) (**Figure 1B**, **Supplementary Data Table 1**, **Methods**). A random sequence (‘dummy’) promoter and the promoter sequence typically used with the transposase and ABR genes were included as internal benchmarks, resulting in 18 total promoter variants per gene.

The same set of promoter variants was cloned upstream of the transposase and the selectable marker, barring internal benchmarks. The overall design of our transposon variant library contains 3888 constructs, comprising of two transposases, six selectable markers, and 18 promoters driving each gene (**Figure 1A**, **Supplementary Data Table 3**). All promoter variants were paired with unique DNA sequence barcodes contained within the transposon. To quantify transposon insertion efficiency, we used the unique DNA barcode combinations to identify each promoter pair and employed transposon insertion sequencing (Tn-seq) to measure their insertion rate in the target genome (**Supplementary Figure 1**).

We first generated six pooled Himar1 transposon variant libraries, one for each of the ABR markers (**Supplementary Figure 2**). Libraries were cloned in *Escherichia coli* and transformed into a conjugative donor strain, *E. coli* BW29427. Amplicon sequencing detected 89.5%-96.9% of the possible 324 combinatorial variants across all six libraries, and long-read Nanopore sequencing verified that libraries were assembled with the correct overall architecture. On average, we detected 306/324 variants per library, with no individual variant comprising more than 1% of the pool (**Supplementary Figure 3**).

We validated performance by conjugating two libraries into readily-transformable strains: *E. coli* DH10B (CAM, SPEC) and *Pseudomonas alcaliphila (i.e., Ectopseudomonas alcaliphila)* (GEN, KAN). Thousands of transconjugants were obtained on selective media, pooled, and analyzed by Tn-seq, identifying 10^3^-10^4^ unique genomic insertion sites across all libraries (**Supplementary Figure 4**). We observed clear strain-specific transposase promoter preferences; for instance, promoter variant 14434 was the most efficient Himar1 promoter variant in *E. coli,* regardless of the antibiotic marker used (**Figure 1C-D**). We also observed ABR marker-specific promoter preferences within strains; for instance, promoter variant 13168 was the top-performing variant in *P. alcaliphila* when using kanamycin but failed to yield any detectable insertion events using gentamicin (**Supplementary Figure 4**). Both strains showed low levels of transposon insertions from variants harboring the dummy promoter, indicating that this random sequence may possess cryptic promoter activity or that transcriptional read-through is occurring from other plasmid regions. We detected no significant correlation between transposition efficiencies observed in our data and the transcription strengths previously published for these promoters in *E. coli* (**Figure 1E-F**). This suggests that transposition efficiency does not depend solely on promoter strength, at least not when all promoters are known to be active in *E. coli a priori*.

### Screening Himar1 transposon variant libraries in 92 bacterial strains

To enable scalable screening of our transposon variant libraries across diverse non-model bacteria, we developed an automated 96-well plate method for parallel testing in multiple strains, which we refer to as the low-input transposon variant screen (**Supplementary Figure 5**). This *E. coli*-based conjugation approach delivers libraries to target strains, after which pooled samples are selected in antibiotic-containing media and growth is measured compared to a negative control (*i.e.*, conjugation with a donor harboring a plasmid with the same ABR gene, but no transposon elements). Samples showing significant growth above the control are analyzed by Tn-seq.

Using this method, we delivered all six Himar1 transposon variant libraries to 92 diverse bacterial strains spanning 12 phyla. We observed positive readouts for 43 (47%) of the recipient strains, with an average of 4 positive samples per recipient, and were able to identify high confidence functional transposons for 14 (33%) of those positive strains (**Figure 2A-B**, **Supplementary Table 2**, **Supplementary Data Table 5-6**). Specifically, these strains had at least one sample sequenced with more than 100 transposon insertion sites (termed ‘high confidence’), a threshold at which the R^2^ of promoter efficiency replicability is greater than 0.5 on average (**Figure 2C**). High-confidence strains span five taxonomic classes: Gammaproteobacteria, Betaproteobacteria, Alphaproteobacteria, Verrucomicrobiae, and Clostridia (**Figure 2A**, **Supplementary Table 2**, **Supplementary Data Table 5-6**). For 29 (67%) of the 43 positive strains, Tn-seq detected only 10-100 insertions; these strains had low transposition efficiency, significant rates of plasmid retention, difficulty in assigning genomic insertion sites with read-mapping, or a combination of these challenges (termed “low confidence”). Of the remaining 49 strains, 40 had no results and 9 had fewer than 10 detectable insertions which we consider failures as it can be challenging to distinguish very few insertion sites from mismapping from plasmid and donor sequences.

**Figure 2.**
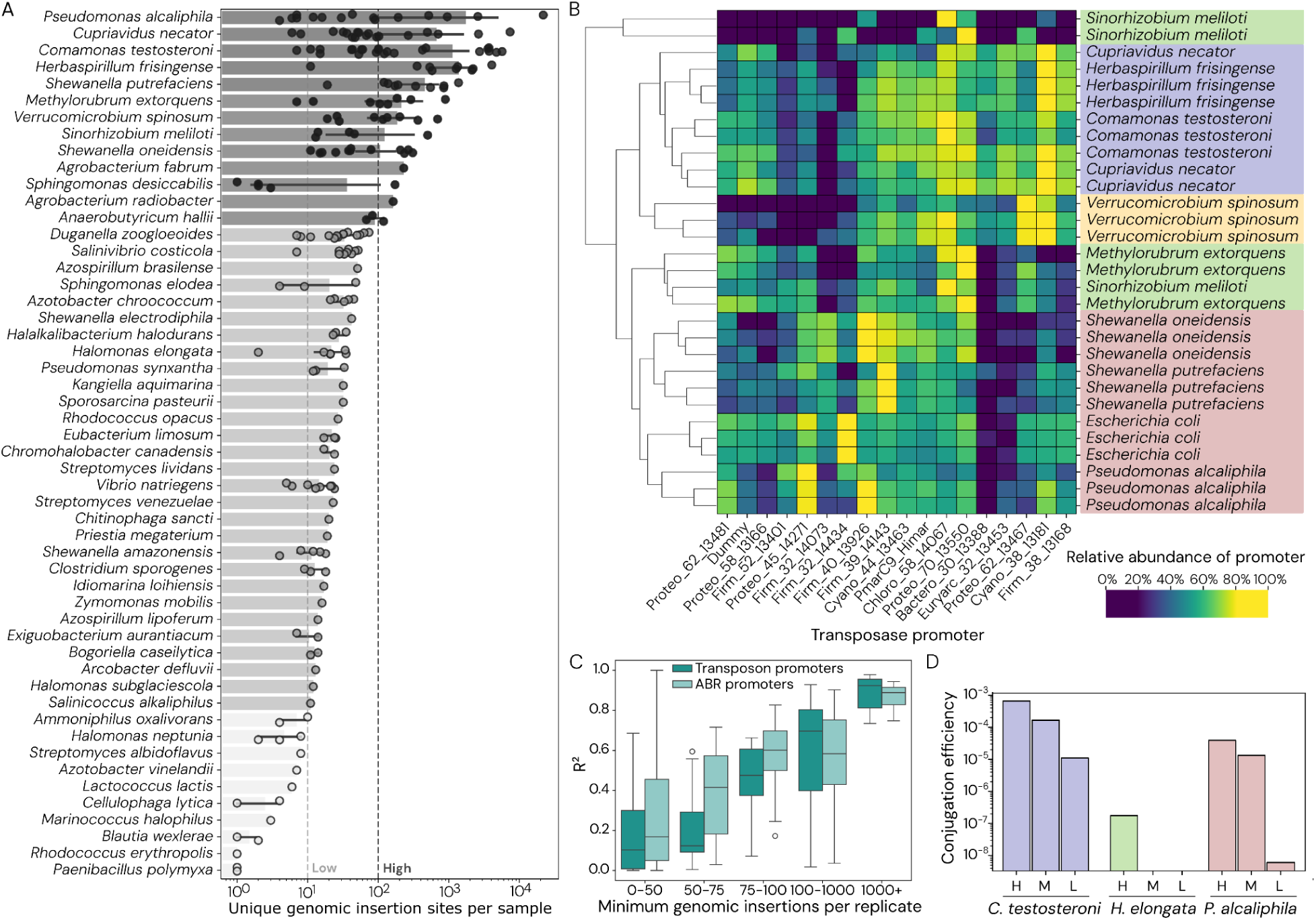
Himar1 transposon variant screen on a cohort of 92 bacterial species. (**A**) Bacterial strains ranked by the number of unique genomic insertion events measured by Tn-seq analysis. ‘High confidence’ strains received over 100 insertions per sample, while ‘low confidence’ strains received between 10-100. Each data point represents the total number of unique genomic insertions sites obtained from delivery of a 324-member transposon variant library. Note the data for *Cupriavidus necator* is a combination of two different strains (see Supplementary Table 2) (**B**) Himar1 transposase promoter efficiencies in strains with at least 100 observed insertion events in multiple samples. Each row represents a single replicate, and rows are clustered by relative transposase promoter efficiencies. Strains are colored by their class: Gammaproteobacteria (red), Alphaproteobacteria (green), Betaproteobacteria (purple), or Verruomicrobia (yellow). (**C**) The replicability of promoter efficiencies as measured by Tn-seq across all microbes screened consistently increases with transformation efficiency for both transposase and ABR promoters. (**D**) Validation of promoter efficiencies. Promoters with high (H), medium (M), and low (L) abundances from the screen results were individually cloned and then tested for their conjugation efficiencies in three strains. Conjugation efficiency is reported as the number of transconjugants per recipient.

Notably, seven of the positive strains in our assay had no published transposon libraries, including the Gammaproteobacteria *P. alcaliphila* (a fast-growing psychrophile and halophile^26^) and *Shewanella putrefaciens* (a metal-reducing marine bacterium^27^), and the Betaproteobacteria *C. testosteroni* (a degrader of plastic- and lignin-derived compounds^28^)*, Herbaspirillum frisingense* (a nitrogen-fixing plant endophyte^29^), *Duganella zoogloeoides* (an exopolysaccharide producer^30^), and *Sphingomonas dessicabilis* (a desiccation-resistant desert soil bacterium^31^) (**Figure 2A-B**). We also observed numerous transposon insertions across multiple samples in the anaerobic human gut commensal *Anaerobutyricum hallii*^32^, a species with no previously reported genetic tools. In addition, we identified a functional transposon construct for *Verrucomicrobium spinosum*, a strain in the divergent Verrucomicrobiota phylum which only comparatively recently had transposon mutagenesis published^33–35^, but this is the first quantification of relative promoter efficiencies in a strain from this phylum. For all strains with high-confidence transposition results, our screen identified promoters with superior efficiency to the endogenous Himar1 transposase promoter on the pMarC9-R6k vector^36^, demonstrating the significance of identifying host-specific promoters to achieve high efficiency transposition in new microbes. On average, the degree of improvement of the best promoter compared to the endogenous transposase or ABR gene promoters ranged from ∼2-20-fold (**Supplementary Figure 6**).

For the bacterial strains with over 100 insertion sites identified in more than one sample, we found that relative promoter efficiencies were strain-specific and reproducible (**Figure 2B**). Further, these promoter preferences often clustered at higher taxonomic ranks. For example, the Betaproteobacteria *Cupriavidus necator*, *H. frisingense*, and *C. testosteroni* consistently preferred similar promoters, while the Gammaproteobacteria *Shewanella oneidensis*, *S. putrefaciens*, *P. alcaliphila*, and *E. coli* preferred an overlapping but distinct set of promoters. This indicates a degree of taxonomic conservation of promoter pair preferences across strains in a given clade, potentially alleviating the need to screen future strains in that clade. For example, promoter variant 1381 was the optimal transposase promoter across all three aforementioned Betaproteobacterial strains, while promoter variant 13550 was optimal in the Alphaproteobacteria *Methylorubrum extorquens* and *Ensifer meliloti* (formerly *Sinorhizobium meliloti*).

To independently validate the results of the transposon variant screen, we individually cloned and tested single constructs with relatively high, medium, or low transposition efficiencies as predicted by their pooled Tn-seq abundances in three stains: *P. alcaliphila*, *Halomonas elongata*, and *C. testosteroni* (**Supplementary Data Table 4, 6**). We found that transposon variants ranked high in our pooled screen produced the highest transposition efficiencies in each strain, while medium-and low-ranked promoters produced correspondingly lower transposition efficiencies (**Figure 2D**).

Transposition efficiencies varied widely across different strains. However, with even 100 insertional events, the replicability of promoter preferences is high, and therefore, this high-throughput screen is still well-suited for identifying optimal transposon variants that can subsequently enable much larger mutagenesis libraries (**Figure 2C**). Further, replicability increases in the screen with insertional efficiency: across all microbes we delivered to, replicate-replicate correlations increased with overall insertional efficiency (**Figure 2C**). In recipients with more than 500 observed insertional events, promoter efficiencies replicated with R^2^ values on average >0.75 (**Figure 2C**).

While we successfully identified functional transposons for 47% (43/92) of tested strains, our assay did not yield results for the remaining strains. Specifically, for 30% (28/92) of strains we observed growth in our assay but few or no genomic insertion sites were identified by Tn-seq, and for the remaining 23% (21/92) of tested strains we observed no transconjugants (*i.e.*, no growth under antibiotic selection).

Several factors likely contribute to these cases. First, the 96-well plate format, while enabling parallel screening of diverse strains, requires reduced donor and recipient biomass compared to standard petri dish conjugations. This increases the limit of detection, potentially missing strains with low transformation efficiency. Second, in many cases sequencing reads predominantly mapped to unmobilized plasmid rather than genomic insertions, likely due to random integration of whole plasmid or persistence of plasmid DNA from the conjugative donor. These cases are excluded as it is difficult to informatically distinguish them from a low number of legitimate genomic insertions (*i.e.*, <20). Third, success rates differed markedly between bacterial groups. While over 50% of Gram-negative strains had transposon insertions, only 29% (12/42) of Gram-positive strains had positive results, with only *Anaerobutyricum hallii* exceeding 100 insertions. This likely reflects lower DNA delivery efficiency from the *E. coli* donor to Gram-positive recipients^37^, which could be improved using alternative delivery methods like electroporation. In addition, Type IV restriction-modification (R-M) systems in recipient genomes were associated with failure to detect transposon insertions (Fisher’s exact test; p=0.006), likely due to the *E. coli* donor’s active *dam* and *dcm* methylases. Using a methylation-deficient conjugative donor or electroporation of unmethylated DNA could address this issue. Finally, our limited set of 16 promoters may be insufficient for effective transposase and ABR gene expression in certain clades. Despite efforts to span diverse phyla, certain clade-specific promoter architectures may not be represented in our design. Based on these findings, transposon variant screening showed highest success rates in Pseudomonadota, with reduced effectiveness expected in other phyla.

Despite these limitations, these data demonstrate that our transposon variant screen can be applied to diverse bacteria in a scalable manner to both establish genetic tractability and identify highly-efficient constructs for transposon mutagenesis in non-model strains.

### Development of a dual-transposase screen

To account for potential variation in the relative efficiencies of different transposases, we next expanded our screen to include both Himar1 and Tn5 transposases. As with the Himar1-only libraries above, we assembled six pooled Tn5-only transposon variant libraries, with one library for each of the ABR markers. We then pooled Himar1 and Tn5 libraries that share the same ABR genes, resulting in a final set of six dual-transposon variant libraries. Sequencing verified the presence of between 89.4-96.8% of the 648 possible variants in each library (**Supplementary Figure 7**). While Tn5 constructs were more abundant than Himar1 constructs on average (62% of the library), variants with both transposases were well represented.

Although previous Tn-seq analysis of Himar1 transposon variant libraries relied on MmeI restriction enzyme sites within the Himar1 mosaic ends, simultaneous Tn-seq analysis of Himar1 and Tn5 transposition necessitated an NGS library preparation method functional with both. Inspired by previous studies^38^, we employed a semi-arbitrary (SemiArb) PCR-based method that simultaneously amplifies the transposon leading end and insertion sites from both Tn5 and Himar1 transposons (**Supplementary Figure 8**).

Using both low-input (96-well-format) and high-input (petri dish-format) conjugations, we delivered the dual-transposon variant libraries to eight strains in which Himar1-only transposon variant libraries previously functioned to various degrees. We detected transposition events driven by both transposase systems in all eight strains with total insertion counts spanning three orders of magnitude (**Figure 3A**). In *Piscinibacter sakaiensis* and *E. coli*, we observed 2-10-fold higher Tn5 transposition efficiency compared to Himar1, while in *C. necator* and *M. extorquens*, the efficiency of Himar1 was 1.1-1.5-fold higher than Tn5 (**Figure 3A**). As expected, for *P. alcaliphila*, the total number of observed insertions increased by one to two orders of magnitude under high-input conditions compared to low-input conditions.

**Figure 3.**
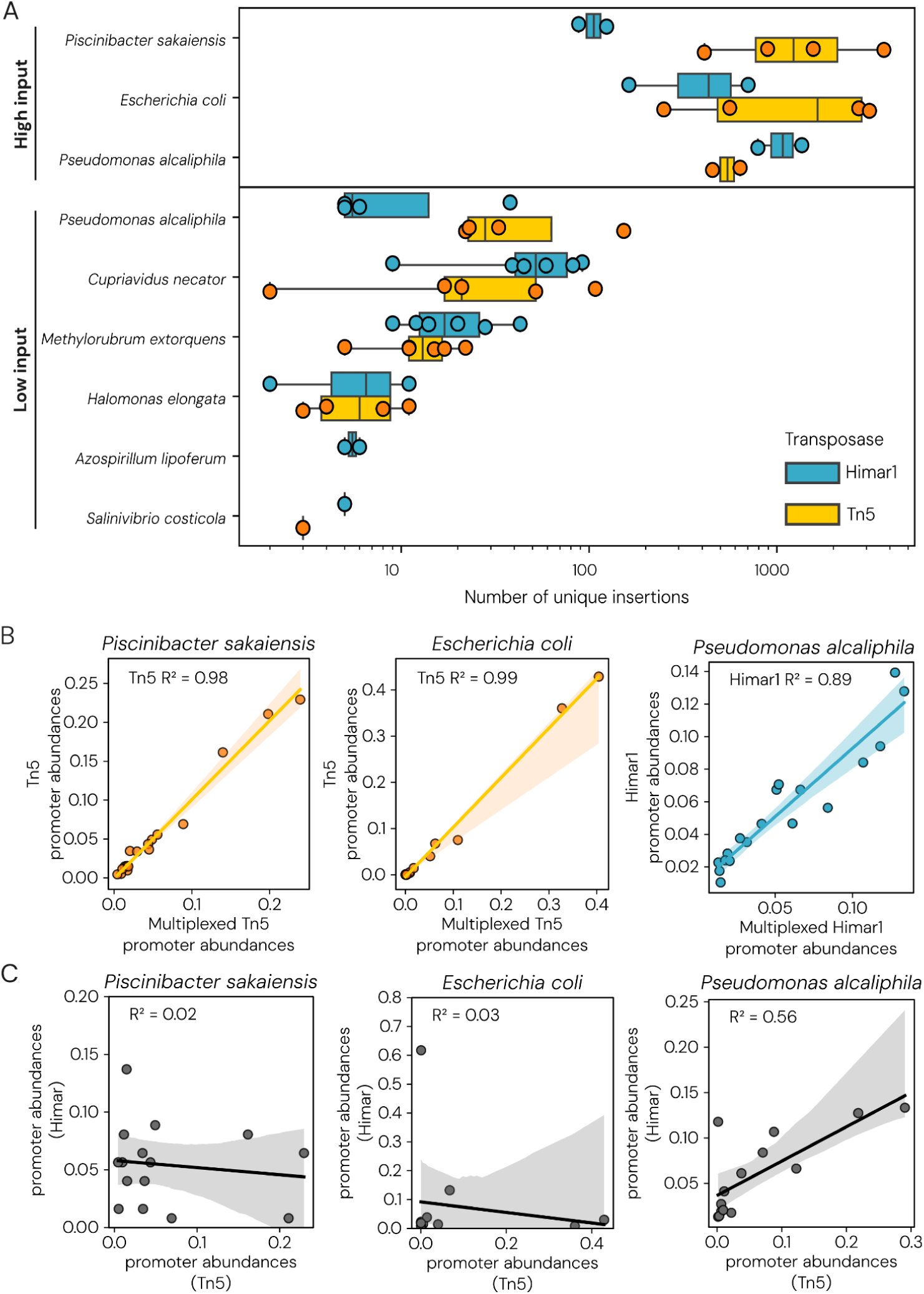
Validation and testing of the multiplexed Himar1 and Tn5 transposon variant screen. (**A**) Number of genomic insertions observed for different recipient strains undergoing dual-transposon promoter screen, as determined by sequencing. Conjugations were performed either under standard petri dish-based format (‘high input’) or 96-well plate-based format (‘low input’) and transconjugants sequenced after growth in selective media. (**B**) Comparison of Tn5 (orange) or Himar1 (blue) promoter abundance between the Himar1-only or Tn5-only transposase library screens (y-axis) and Himar-Tn5 dual-transposase library screen (x-axis). (**C**) Comparing the efficiencies of promoters associated with Himar1 (y-axis) versus Tn5 (x-axis) transposons for three strains.

The observed transposition efficiency of promoter variants for each transposase showed a strong correlation between the dual-transposon variant screen and equivalent single-transposon variant screens (**Figure 3B**). As observed in the Himar1-only transposon variant screen, the transposition efficiencies of promoter variants were highly replicable within strains, especially strains with many (*i.e.*, >100) insertions per sample (**Supplementary Figure 9**). Interestingly, we observed substantial differences between Himar1 and Tn5 transposition efficiencies when driven by the same promoter, even in the same strain. For example, there was no correlation between Himar1 and Tn5 transposition efficiencies when driven by the same promoter in *E. coli*, *P. sakaiensis*, and only a moderate correlation (R^2^=0.56) in *P. alcaliphila* (**Figure 3C**). In other words, in a given strain, the best promoters for the Himar1 transposase are not necessarily the best for the Tn5 transposase, nor the best for the ABR marker. One biological explanation is that overly high expression can cause misfolding, premature transposition, or toxicity of either the transposase or the ABR markers, and that this may happen in a gene-specific manner.

Taken together, these data show that optimal transposition is governed by a complex combination of genomic and cellular factors, highlighting the value of our approach.

### Generation of genome-wide transposon library in *Comamonas testosteroni*

To demonstrate application of our screen for assessing gene function in a new host, we constructed a large randomly-barcoded transposon library in *C. testosteroni* KF-1, a bacterium capable of bioprocessing aromatic carbon compounds such as lignin and plastic breakdown products^39^.

Specifically, our screen identified the top-performing transposon construct as the Himar1 transposase driven by promoter varant 14067 and KAN ABR gene driven by promoter variant 14271 (**Supplementary Data Table 6**, **Supplementary Figure 10**). We then cloned the top performing construct to be used for genome-wide mutagenesis. To facilitate downstream mutant library screening experiments (*i.e.* enable RB-TnSeq^40^), we added a 20 bp random barcode sequence to the transposon.

The barcoded transposon construct was delivered to *C. testosteroni* by conjugation, resulting in a high-efficiency transposon mutant library of ∼7.7 x 10^5^ transconjugants. Using Tn-seq, we detected 230,573 uniquely-barcoded strains with 73,195 unique transposon insertion sites in the genome of *C. testosteroni*. We found that 92% of genes in the genome were represented by at least one insertion in the middle 80% of the gene. There was minimal evidence of genome-wide bias in insertion site frequency (**Supplementary Figure 11**). Therefore, using the top-performing transposon variant construct identified by our screen enabled the creation of a near-saturation transposon mutant library in this strain.

*C. testosteroni* KF-1 has previously been studied for its ability to metabolize terephthalic acid (TER), a constituent monomer of polyethylene terephthalate (PET) plastic, and 4-hydroxybenzoate (4HB), a lignin-decorating moiety^39^. RNA-seq and genomic homolog analyses have generated hypotheses about the genes involved in this degradation pathway^39^, but for almost all of these genes, no functional genetic evidence for their roles is available. To identify genes involved in the degradation of TER and 4HB, we grew the *C. testosteroni* transposon mutant library in minimal media supplemented with TER, 4HB, or succinate (control) as a carbon source, extracted genomic DNA, and performed barcode sequencing (BarSeq) to quantify transposon mutant abundance (**Figure 4A**).

**Figure 4.**
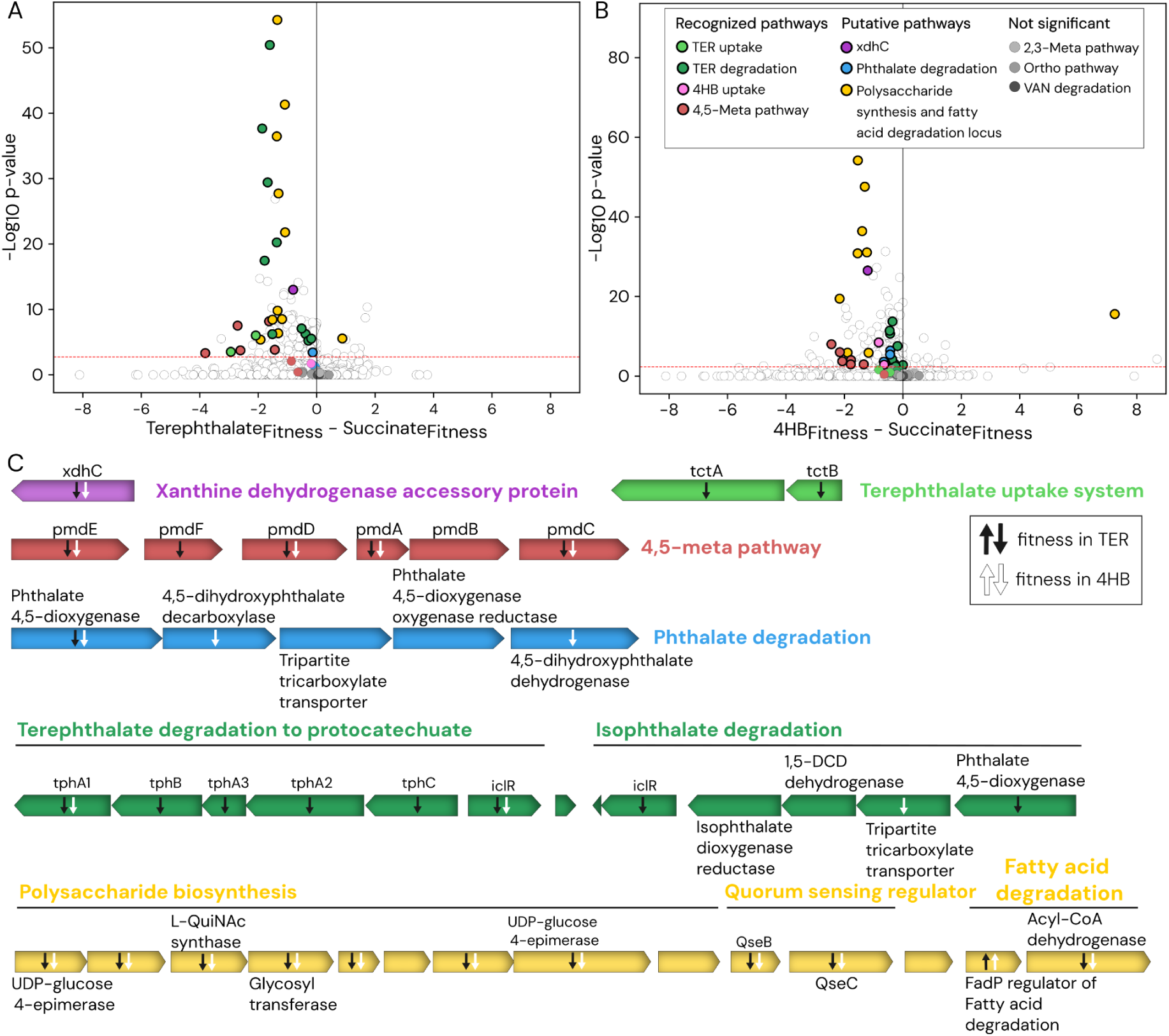
Differential fitness of genes during growth of *C. testosteroni* on terephthalic acid (TER) and 4-hydroxybenzoate (4HB) substrates. (**A**) Volcano plot of gene fitness during growth in TER. The x-axis reflects the degree of the change in fitness (measured by the Log-2 fold change since baseline), and the y-axis reflects the p-value significance. Genes with negative fitness values have improved growth on TER, and genes with positive fitness values have decreased growth on TER. The red line is at a BH-adjusted p-value cutoff of *q*=0.05. (**B**) An equivalent volcano plot for growth in 4HB. (**C**) Genetic loci that contain genes with differential fitness. The operons are colored by pathway, and the colors for each pathway or operon are equivalent to the colors in (A) and (B). Arrows within genes indicate significant negative fitness (downward pointing) or positive fitness (upward pointing) in TER (black) or 4HB (white).

Several genes with known or clearly interpretable functional annotations were found to have significant fitness benefits during growth on TER or 4HB (**Figure 4A**, **Supplementary Tables 7-8**). This experiment provided functional evidence that TER degradation occurs via *tctA* and *tctB* transporters, and conversion to PCA occurs via the *tph* operon, as had been previously predicted through RNA-seq data^39^. These results further confirmed that uptake of 4HB occurs via the *pobA* transporter^39^. They also confirmed that the 4,5-meta pathway, and not the “Ortho” or “2,3-meta” pathways, is the mechanism by which *C. testosteroni* metabolizes the protocatechuate intermediate of both TER and 4HB (**Figure 4A-C**).

The experiment suggests a number of new hypotheses for genes with fitness benefits during growth on TER or 4HB, that were previously unrecognized in C. *testosteroni* KF-1. The fitness screen identified a homolog of a gene previously known to have specificity for phthalate over terephthalate degradation^28^, but which may also play a role in terephthalate degradation in *C. testosteroni* KF-1 (**Figure 4C**, **Supplementary Tables 7-8**). Further, we identified an operon of genes related to polysaccharide biosynthesis, biofilm biosynthesis, and/or quorum sensing as beneficial for growth on both TER and 4HB (**Figure 4A-C**). Finally, we identified one gene where transposon knockout was associated with improved growth on both 4HB and TER: a homolog of the FadP regulator of fatty acid degradation (**Figure 4B-C**). This gene is adjacent to the polysaccharide biosynthesis beneficial for growth on 4HB and TER, and therefore, it may also be a negative regulator of these genes.

### Conclusion

Here, we describe a scalable system for transposon mutagenesis in non-model bacteria. Using this pooled and combinatorial screening workflow, we generated a large and comprehensive dataset characterizing the performance of transposon variants across a wide range of bacterial phyla, and identified functional transposons in 43 non-model bacteria. We demonstrate that top-performing transposons from this approach can be used in large-scale transposon mutagenesis by generating a genome-wide transposon mutant library in *C. testosteroni* for the first time.

Further work is warranted to identify functional transposons in non-model bacteria. Additional considerations include DNA delivery method, host genetic defense systems, and molecular barriers for expression of heterologous DNA (*e.g.*, codon usage or RBS preferences affecting transcription and translation). The described transposon system is scalable with further addition of curation of transposases, promoters, or other regulatory elements, and is another addition to our growing collection of tools to study and engineer non-model organisms.

## Materials and Methods

### Strains and growth conditions

*Escherichia coli* strains were grown in LB medium at 37°C, unless otherwise specified. *E. coli* BW29427 (sourced from B.L. Wanner) cultures were supplemented with 60 µg/mL diaminopimelic acid (DAP). *Comamonas testosteroni* KF-1 received from the Leibniz Institute DSMZ, Germany (DSM 14576) was grown in NB medium at 30°C, unless otherwise specified. All bacterial recipient strains and their growth, conjugation, and selection media are summarized in **Supplementary Data Table 5**. Solid media plates were prepared with 1.5% agar and liquid cultures were grown with shaking at 225 RPM. Standard antibiotic concentrations used for selection were 100 µg/mL carbenicillin (CARB), 50 µg/mL kanamycin (KAN), 20 µg/mL gentamicin (GEN), 4 µg/mL erythromycin (ERM), 10 µg/mL tetracycline (TET), 34 µg/mL chloramphenicol (CAM), and 50 µg/mL spectinomycin (SPEC). Antibiotic supplementation outside of this standard concentration is noted where relevant.

### General molecular biology techniques

Genomic DNA (gDNA) extractions were performed either using the DNeasy Blood & Tissue Kit (Qiagen 69506) or the Applied Biosystems MagMAX™ Viral/Pathogen Ultra Nucleic Acid Isolation Kit (Thermo Fisher Scientific A42356) on the KingFisher (Thermo Fisher), with the latter used for all transposon variant screen DNA extractions. Concentrations of nucleic acids were determined using a Qubit Flex Fluorometer (Fisher Scientific Q45893) using the Qubit™ 1X dsDNA High Sensitivity Kit (Thermo Fisher Scientific Q33231). Plasmid DNA extraction from *E. coli* was performed using the QIAprep Spin Miniprep Kit (Qiagen 27106).

### Transposon variant library design

The transposon variant libraries described in this study consist of 18 separate pooled plasmid libraries, each composed of 324 unique plasmid constructs. Each library was generated from a single destination vector into which 18 promoter variants driving transposase expression and 18 promoter variants driving antibiotic resistance (ABR) marker gene expression were assembled in a pooled format (**Supplementary Data Table 1-3**).

Sixteen promoter variants were selected by iteratively filtering the 421 promoters from Yim et al. 2019. Promoter sequences with BsaI cut sites were removed from the set. To choose promoters more likely to be avoided by recipient R-M systems, promoter sequences were scanned for all palindromic methylated motifs reported in the REBASE database of restriction-modification systems^41^. Promoters with more than a 15% probability of being targeted by restriction enzymes across species in the dataset were then removed from consideration. Constrained K-means clustering was then used with parameters n_clusters=4, size_min=50, size_max=150, to identify clusters of relative transcriptional strength measured in 10 bacteria as reported by Yim et al. 2019. For each transcriptional strength cluster, promoters were ranked by GC content and one promoter was selected from each 25% quantile of GC content, preferentially selecting promoters that originated from a unique phylum, or if no unique phylum was present in the bin, choosing the promoter closest to the mean.

In addition to these 16 promoter variants, we designed positive and negative control promoter variants to be included as internal controls within transposon variant libraries. For positive control promoter variants, we selected the native/standard promoter used in the literature for the downstream gene of interest: transposase enzyme or ABR gene. For negative control promoter variants, we generated 165 bp random DNA sequences with ∼50% GC content (**Supplementary Data Table 1**).

To format transposase promoter variants for library construction, the relevant transposon mosaic end sequence was first added upstream of the promoter sequence. Next, a constant 30 bp primer binding sequence was added upstream of the mosaic end and a unique 20 bp DNA barcode sequence was added upstream of that primer binding sequence. Finally, sequences were flanked with inward-orientated BsaI restriction enzyme cut sites to enable downstream Golden Gate assembly reactions. For ABR gene promoter variants, another 20 bp constant primer binding region was added upstream of the promoter sequence and a unique 20 bp DNA barcode sequence was added upstream of that primer binding sequence. Finally, sequences were again flanked with inward-orientated BsaI restriction enzyme cut sites to enable downstream Golden Gate assembly reactions (**Supplementary Figure 2**). This entire set of sequences was commercially synthesized as sequence-verified parts within a plasmid (Twist Biosciences and Integrated DNA Technologies). However, in one case, pGLU_119, commercial plasmid construction failed, and therefore in-house cloning from commercial DNA fragment synthesis was performed.

We then designed a set of 12 destination vectors for library cloning via Golden Gate assembly (**Supplementary Data Table 2**). Each of these destination vectors shares several common features: an RK2/RP4 origin of transfer (*oriT*) sequence for conjugative delivery, the conditional *E. coli* origin of replication R6Kγ, an *E. coli* ABR marker (CARB), a transposon mosaic end sequence, and a Golden Gate dropout part consisting of an mScarlet reporter with constitutive expression in *E. coli* flanked by inward-oriented BsaI restriction enzyme cut sites to enable downstream Golden Gate assembly reactions. The latter dropout part itself is flanked by a promoter-less ABR gene marker and a promoter-less transposase gene variant (two possible variants). The 12 destination vectors differ in the identity of the transposase system (Himar1^36^ or Tn5^24^) and ABR gene marker (KAN, GEN, ERM, TET, CAM or SPEC^20^). The Golden Gate assembly cut sites create overhangs that allow the introduction of promoter variants from the previous section in such a way that enables expression of the ABR gene and transposase gene coding sequences. Each of these 12 plasmids was obtained using commercial synthesis of sequence-verified plasmids (Twist Biosciences).

All transposase promoter variants, ABR gene promoter variants, and destination vectors are being made available to order via Addgene as both individual plasmids and as a single toolkit with all plasmids, “Cultivarium JERBOA Toolkit”.

### Transposon variant library construction

First, six separate ABR gene promoter pools (pGLU_35-40) were made by manually mixing equal amounts of 18 different promoter plasmids (each 250 nM) (**Supplementary Data Table 3**). The six promoter libraries differ only in the identity of the ‘positive control’ promoter specific to each of the six ABR genes. Next, two transposase promoter pools were made by manually mixing equal amounts of 18 different transposase promoter plasmids (each 250 nM) (**Supplementary Data Table 3**). Since these promoter sequences contain a specific transposon mosaic end sequence, two separate pools were required for the two distinct transposon systems (Himar1 and Tn5). In addition, the two libraries differ in the identity of the ‘positive control’ promoter specific to each of the two transposases used.

To create separate Himar1 and Tn5 transposon variant libraries, 12 BsaI Golden Gate assembly reactions were performed, each using the appropriate combination of destination vector, ABR gene promoter pool, and transposase gene promoter pool (**Supplementary Data Table 3**). Specifically, in each reaction, 3 μL of the destination vector (20 nM stock, providing ∼60 fmol per reaction) was combined with 1 μL of each pooled promoter library (for Himar1 transposon variant libraries: 250 nM stock, ∼250 fmol per reaction; for Tn5 transposon variant libraries: 20 nM stock, ∼20 fmol per reaction). A master mix was prepared on ice with the following components: 1 µL nuclease-free molecular biology grade water (MGW), 3.33 µL NEBridge Ligase Master Mix (New England Biolabs M1100S, 3X), and 0.67 µL BsaI (New England Biolabs R3733L, 20 U/µL) per reaction. The total volume of the master mix was scaled according to the number of reactions, and 5 µL of the master mix was added to each reaction tube. Reaction mixtures were mixed thoroughly by pipetting and briefly centrifuged using a mini-spin centrifuge. Reactions were processed on a BioRad thermocycler using NEB’s recommended cycling conditions for NEBridge Ligase Master Mix Golden Gate assembly with BsaI. Following the assembly reaction and a 5 minute 60°C incubation to digest away any remaining destination vector DNA, the resulting constructs were purified using the DNA Clean & Concentrator-5 kit (ZYMO D4013), eluted in 6 µL, and transformed into *E. coli* BW29427 electrocompetent cells by electroporating with the following condition: 2.0 kV, 200 Ω, and 25 μF. Cells were recovered in SOC supplemented with 60 µg/mL DAP at 37°C for one hour, then added to 100 mL of LB medium supplemented with 60 µg/mL DAP and 100 µg/mL CARB. Cells were grown overnight, a 5 mL aliquot was harvested for plasmid extraction and downstream variable promoter barcode sequencing (VBBarSeq) and the remainder of the culture was cryo-stocked in 1 mL aliquots supplemented with glycerol to a 15% (w/v) final concentration. VBBarSeq sequencing to characterize the assembled libraries was performed using primers oCG0038 and oCG0039 (**Supplelementary Table 4**) according to the amplicon-sequencing protocol described in Gilbert et al. 2023^9^.

Finally, to create a combined Himar1 and Tn5 transposon variant library, 1 mL aliquots of individual transposon variant libraries from the previous section were inoculated into 50 mL liquid cultures and grown until an OD_600_ of 0.4-0.8 was reached. Six pools were then prepared by mixing OD-normalized volumes of Himar1 and Tn5 libraries, pairing the 12 libraries by shared ABR genes. Again, 5 mL aliquots were harvested for plasmid extraction and downstream variable promoter barcode sequencing (VBBarSeq) and the remainder of the culture was cryo-stocked in 1 mL aliquots supplemented with glycerol to a 15% (w/v) final concentration.

### Transposon variant screen

Initial Himar1-only library validations (**Figure 1C-D**, **Supplementary Figure 4**), validation of individual transposon variant performance (**Figure 2D**), *P. sakaiensis*, *E. coli* and *P. alcaliphila* dual-transposon variant screen experiments (**Figure 3A**), and *P. sakaiensis* and *E. coli* Tn5-only transposon variant screen experiments (**Figure 3B**) were all performed with the high-input protocol. Unless otherwise specified in **Supplementary Table 2**, screening of strains in the non-model cohort transposon variant screen (**Figure 2A**) and of the remaining dual-transposon variant screen strains (**Figure 3A**) were performed with the low-input protocol.

#### High-input protocol

Conjugation was performed as previously described^20^ with the following modifications. *E. coli* BW29427 conjugative donor harboring either plasmid libraries or single control plasmids (**Supplementary Table 3**) were each grown in 50 mL of LB medium supplemented with CARB (libraries) or CAM (control plasmids). Recipient strains were cultured in 50 mL of their respective recommended growth media (**Supplementary Data Table 5**). Donor and recipient cultures were grown for 2 hours, adjusted to an OD_600_ of 100, and mixed at a 1:1 ratio (by volume, 50 µL of each) on a conjugation plate (petri dish). LB 1.5% agar medium supplemented with 60 µg/mL DAP was used as the conjugation media. Conjugation plates were incubated for 20 hours at 30°C, unless otherwise stated. After scraping the conjugation mixture off conjugation plates, the suspensions were washed by centrifuging at 5000 RCF for 10 min at room temperature, resuspending in 1 mL of PBS, centrifuging again, and finally resuspending in 500 µL of PBS. Note, this wash step was not completed for *P. sakaiensis* as cells did not pellet well post-conjugation. Transconjugants were selected on recipient recommended media 1.5% agar plates supplemented with the appropriate antibiotic using standard concentrations, unless otherwise stated. Colonies were counted manually after 1-4 days.

#### Low-input protocol

All liquid transfers (supernatant removal, resuspension, mixing, etc.) were performed using the Opentrons Flex, unless otherwise specified. Up to eight recipient strains and the six control donors (**Supplementary Table 3**) were streaked onto 1.5% agar plates of the recommended media from glycerol stocks and incubated at 37°C (donors) or the recipient optimal growth temperature until colonies appeared. Control donors were inoculated into 1 mL of LB supplemented with 60 µg/mL DAP and 20 µg/mL CAM in the even-numbered top wells of a 24-well column reservoir. For the library donors, one aliquot of each *E. coli* BW29427 transposon variant library glycerol stock (pGLU_28-33) was thawed at room temperature and then transferred into the odd-numbered top wells of the same 24-well column reservoir. Then, 980 µL of LB supplemented with 60 µg/mL DAP and 100 µg/mL CARB (library donors) or 20 µg/mL CAM (control donors) was dispensed into each well of a 96-deepwell plate. Then, 20 µL of each library and control donor in the 24-well column reservoir were used to inoculate the deepwell plate, keeping the same donor in each column, pipetting down all the rows (**Supplementary Figure 5**). Recipient strain cultures were also set up in a 96-deepwell plate row-wise, keeping the same recipient in each row, pipetting across all the columns. 980 µL of recipient recommended media was first added to the appropriate wells before being inoculated with 20 µL of biomass from the streak plate resuspended in 1 mL media (until cloudy). Both deepwell plates were sealed with a breathable cloth membrane and then incubated at 30°C (donors) or recommended temperature, shaking at 800 RPM for 20 hours. OD_600_ measurements were taken for both donor and recipient cultures to check for growth. If there was no growth, a suspension prepared directly from the fresh streak plate was used instead.

Conjugations were performed in 96-deepwell plates containing 200 µL of LB 1.5% agar supplemented with 60 µg/mL DAP that were dried for 1 hour under a laminar flow hood or similar. Donors were pelleted by centrifugation at 5,000 RCF for 10 minutes at 4°C, 900 µL of supernatant was removed, and cells were resuspended in 900 µL PBS. Then, both donors and recipients were pelleted at 5,000 RCF for 10 minutes at 4°C, 900 µL of supernatant was removed, and cells were resuspended in 100 µL of the remaining supernatant. Recipient suspensions were then transferred to the donor 96-deepwell plate, mixed, and 20 µL of the conjugation mix then transferred to the LB+DAP agar plate. Conjugations were sealed with a breathable cloth membrane and incubated at the optimal temperature for recipient growth (generally 30°C) for 20 hours. After incubation, 100 µL PBS was added to each well and the plate was shaken at 800 RPM for 5 minutes to resuspend cells off the agar surface. An additional 400 µL PBS was added to each well to dilute the recovered conjugation volume before taking 50 µL out to inoculate 950 µL cultures in both non-selective (recommended recipient media only) and selective (recommended recipient media and appropriate antibiotic supplement according to library donor) media 96-deepwell plates; three or four selective plates were prepared with increasing concentrations of the relevant antibiotics. 100 µL was taken from the non-selective plates to determine the starting OD_600_. Outgrowth plates were sealed with a breathable cloth membrane and incubated at the optimal temperature for recipient growth (generally 30°C) for 24-72 hours, or potentially longer depending on the strain. OD_600_ readings were taken at 24 hours, 48 hours, and beyond as needed to check for growth. At each of these timepoints, samples that had detectable growth in their selective well with an OD_600_ reading at least 0.25 above the corresponding non-selective well were regarded as hits and collected for gDNA extraction by spinning at 13,000 RPM for 3 minutes at room temperature, removing the supernatant, and freezing the cell pellet at −80°C.

### Cloning individual transposon variant plasmids

Individual, clonal transposon variant plasmids were constructed following transposon variant screening. To generate specific constructs, the relevant destination vectors, transposase promoter plasmids, and ABR gene promoter plasmids were first selected. These were then combined via BsaI Golden Gate assembly using NEBridge Ligase Master Mix according to the protocols described in Gilbert et al. 2024. Details of all Golden Gate assembly reactions performed are listed in **Supplementary Data Table 4**. All constructs were sequence-verified by whole plasmid sequencing (Plasmidsaurus).

### Variant barcode transposon insertion sequencing

Two different library preparations were used for the variant barcode Tn-seq of transposon variant screen samples. The MmeI Tn-seq libraries (MmeI VB-TnSeq) can only be used with Himar1 transposon mutants, while the semi-arbitrary PCR libraries (SemiArb VB-TnSeq, **Supplementary Figure 7**) can be used with both Himar1 and Tn5 transposon mutants. gDNA extraction of transposon variant samples was mostly done with the Applied Biosystems MagMAX™ Viral/Pathogen Ultra Nucleic Acid Isolation KingFisher Kit, with some smaller experiments processed with the DNeasy Blood & Tissue Kit.

#### Preparation of MmeI VB-TnSeq libraries

A 30 µL MmeI (New England Biolabs R0637, 2 U/µL) restriction enzyme digest reaction in 1X rCutSmart Buffer (New England Biolabs B6004, 10X) was prepared with 3.6 U of enzyme and 1 µg of input gDNA. Reactions were incubated at 37°C for 15 minutes. For Antarctic Phosphatase (AnP, New England Biolabs M0289) dephosphorylation post-digestion, a master mix of 4 µL AnP Reaction Buffer (10X), 2 µL AnP (5 U/µL), and 4 µL MGW was added directly to each sample and incubated at 37°C for 30 minutes, followed by a 20 minute heat inactivation at 70°C. DNA fragments were then purified using AMPure XP Beads (Beckman Coulter A63881) according to the manufacturer’s protocol using a 1.8X bead to sample volume ratio and eluted in 10.5 µL MGW. The eluted DNA was transferred to a fresh tube for 5 µL IDT pMarC9 P5 adapter duplex (1 µM, **Supplementary Table 4**) and 15 µL Instant Sticky Ligase Mix (New England Biolabs M0370, 2X) to be added in the next ligation step. After incubation at room temperature for 5 minutes, the ligations were purified using the same 1.8X AMPure XP bead cleanup as described above, except with a 13 µL elution volume.

The eluted DNA was then transferred to a fresh tube for the first PCR amplification step with KAPA Hifi HotStart PCR Kit (Roche 07958897001) and EvaGreen Dye (Biotium 31000). A 20 µL reaction was prepared by adding 4 µL KAPA Hifi Fidelity Buffer (5X), 0.6 µL KAPA dNTPs (10 mM), 0.4 µL KAPA HiFi Polymerase (1 U/µL), 1 µL EvaGreen dye (20X), 0.5 µL i5xx primer (10 µM), and 1 µL oCG0039 primer (10 µM) to the eluted DNA. The reaction was completed in a qTOWER^3^G thermal cycler (Analytik Jena) using a standard thermal cycling protocol according to the manufacturer’s instructions: 65°C annealing temperature and 15 second extension time. The progress of each sample was tracked through the amplification curves produced in the qPCR real-time, and the reaction was halted upon seeing sure signs of amplification above the reaction’s baseline - typically around 12-15 cycles.

Next, 1 µL of a 10-fold dilution of the first PCR product was used as template for the second PCR amplification. Reactions were performed using the KAPA HiFi HotStart PCR Kit with EvaGreen Dye in a 20 µL qPCR reaction, following the manufacturer’s protocol and using 1 µL EvaGreen dye (20X), 0.6 µL P5 primer (10 µM), and 0.6 µL i7xx primer (10 µM). The reaction was completed in a qTOWER^3^G thermal cycler using a standard thermal cycling protocol according to the manufacturer’s instructions: 68°C annealing temperature and 15 second extension time. The progress of each sample was tracked through the amplification curves produced in the qPCR real-time, and the reaction was halted upon reaching the stage of exponential amplification - typically around an Intensity [I] value of 50000 RFU and generally around 15 cycles. Products were visualized on an Invitrogen E-Gel EX 2% Agarose gel (Invitrogen G402022) to confirm library size of 288 bp and purity. A double-sided AMPure XP bead cleanup was performed according to the manufacturer’s protocol using an initial 0.4X bead to sample volume ratio and 0.8X final total bead to sample volume ratio to enrich for DNA fragments between 200-300 bp. Cleaned libraries were eluted in 12.5 µL 10 mM Tris-HCl (pH 8.0) with 0.1 mM EDTA.

#### Preparation of SemiArb VB-TnSeq libraries

The method is depicted in **Supplementary Figure 8**. An arbitrary priming reaction was done in a 20 μL volume using the KAPA HiFi HotStart PCR Kit with a recipe slightly modified from the manufacturer’s protocol: 4 μL KAPA HiFi Fidelity Buffer (5X), 0.6 μL KAPA dNTP Mix (10 mM), 1 μL JL0075 primer (50 μM, **Supplementary Table 4**), 0.8 μL KAPA HiFi HotStart DNA Polymerase (1 U/μL), and the remaining 13.6 μL volume split between the appropriate template DNA volume for 500 ng total input and MGW. The reaction was completed in a qTOWER^3^G thermal cycler using the following thermal protocol: 95°C for 5 minutes and 5 cycles of 98°C denaturation for 20 seconds, 23°C annealing for 30 seconds, 68°C extension for 15 seconds. The entire reaction was then purified through a double-sided SPRIselect (Beckman Coulter B23318) bead clean according to the manufacturer’s protocol using an initial 0.5X bead to sample volume ratio to exclude large fragments above 800 bp, followed by taking the supernatant and adding beads for a 1X final total bead to sample volume ratio to exclude small fragments below 250 bp. The bead cleaned product was resuspended in 12.5 μL MGW.

The eluted DNA was then transferred to a fresh tube for the first PCR amplification step with the KAPA Hifi HotStart PCR Kit and 20X EvaGreen Dye. A 20 µL reaction was prepared according the manufacturer’s protocols using 4 μL KAPA HiFi Fidelity Buffer (5X), 0.6 μL KAPA dNTP Mix (10 mM), 0.5 μL JL0076 primer (10 μM), 1 μL JL0073 primer (10 μM), 0.4 μL KAPA HiFi HotStart DNA Polymerase (1 U/μL), and 1 μL EvaGreen Dye (20X). The reaction was completed in a qTOWER^3^G thermal cycler using a standard thermal cycling protocol according to manufacturer’s instructions with a 63°C annealing temperature and 30 second extension time. The progress of each sample was tracked through the amplification curves produced in the qPCR real-time, and the reaction was halted upon seeing sure signs of amplification above the reaction’s baseline - typically around 15-18 cycles. The entire reaction was then purified through a double-sided SPRIselect bead clean according to the manufacturer’s protocol using an initial 0.45X bead to sample volume ratio to exclude large fragments above 900 bp, followed by taking the supernatant and adding beads for a 0.9X final total bead to sample volume ratio to exclude small fragments below 275 bp. The bead cleaned product was resuspended in 20 μL MGW.

Next, 1 µL of a 20-fold dilution of the first PCR product was added to the second PCR amplification as template for another standard KAPA HiFi HotStart PCR Kit with EvaGreen Dye qPCR reaction done in a 20 µL volume exactly according to the manufacturer’s protocol using 0.6 μL i5xx primer (10 μM), 0.6 μL JL0048 primer (10 μM), and 1 µL EvaGreen dye (20X). The reaction was completed in a qTOWER^3^G thermal cycler using a standard thermal cycling protocol according to manufacturer’s instructions with a 71°C annealing temperature and 30 second extension time. The reaction was again halted upon seeing sure signs of amplification above the reaction’s baseline - typically around 14 cycles.

Finally, 1 µL of a 20-fold dilution of the second PCR product was added to the second PCR amplification as template for another standard KAPA HiFi HotStart PCR Kit with EvaGreen Dye qPCR reaction done in a 20 µL volume exactly according to the manufacturer’s protocol using 0.6 μL P5 primer (10 μM), 0.6 μL i7xx primer (10 μM), and 1 µL EvaGreen dye (20X). The reaction was completed in a qTOWER^3^G thermal cycler using a standard thermal cycling protocol according to manufacturer’s instructions with a 68°C annealing temperature and 30 second extension time. The progress of each sample was monitored in real time by qPCR, and the reactions were halted upon reaching the exponential phase of amplification, typically corresponding to an Intensity [I] value of approximately 50,000 after 10-12 cycles. The entire reaction was then purified through a double-sided SPRIselect bead clean according to the manufacturer’s protocol using an initial 0.4X bead to sample volume ratio to exclude large fragments above 1500 bp, followed by taking the supernatant and adding beads for a 0.8X final total bead to sample volume ratio to exclude small fragments below 300 bp.

#### Sequencing of VB-TnSeq libraries

The purified library was quantified using the Qubit™ 1X dsDNA High Sensitivity Kit. The rest of the library preparation was completed according to the Denature and Dilute Libraries Guide for MiniSeq System by Illumina, then loaded and sequenced on an Illumina MiniSeq in a 2 x 150 bp paired-end run, generally targeting at least 100,000 reads per individual sample library.

### Variant barcode transposon insertion library analysis

Quality and adapter trimming of reads was performed using BBDuk from the BBTools^42^ package v39.33 with the following settings: k=19, qtrim=r, trimq=10, maq=10, entropy=0.3. Cutadapt v5.1^43^ was used to extract genomic insertion sequences from forward reads with the settings: -a taacaggttggatgataagtccccggtcta, --minimum-length 10, --overlap 10, --maximum-length 30, --error-rate 10. Variant promoter barcodes were extracted from reverse reads using Cutadapt v5.1 with the following settings: -g CCAGCTTCACACGGCGTGTGGCTGCGGAAC, --minimum-length 30, --overlap 10, --error-rate 10. Genomic reads corresponding to all possible plasmid sequences were identified and excluded using the VSEARCH^44^ --usearch_global command with the following settings: --query_cov 0.75, --id 0.9, --minseqlength 10. Genomic sequences were then mapped to the assembled genomes for each strain using bowtie2 v2.5.4^45^ with default parameters. A custom script with the Pysam library (see Data and Resource Availability) was used to filter mapped reads with a MAPQ score >20 (indicating a strongly unique mapped position) and with ≥90% identity to the reference. Barcode pairs were mapped to the database of promoter barcodes using the VSEARCH --usearch_global command with the parameters: --id 0.9, --target_cov 0.9. The barcode combination supported by the most reads at a given unique genomic site was counted, retaining only barcode combinations supported by at least 3 read pairs.

### Generation of a *C. testosteroni* genome-wide mutant library

The best-performing transposon construct from the transposon variant screen was used to generate a genome-wide mutant library in *C. testosteroni*.

#### Construction of a barcoded transposon library

Using pGLU_127 as a template, barcodes were added using N-mers from IDT primers (**Supplementary Table 4**) to make the pGLU_127_RB barcoded plasmid library. This PCR was a standard KAPA HiFi HotStart PCR Kit reaction done in two separate 50 µL volumes according to the manufacturer’s protocol using JL0061 and JL0062 primers and 1.25 ng input template. The reaction was completed in a qTOWER^3^G thermal cycler using a standard thermal cycling protocol according to the manufacturer’s instructions: 66°C annealing temperature and 3 minute 45 second extension time for 20 cycles. The PCR product was cleaned with the DNA Clean & Concentrator-5 kit, eluting in 25 µL MGW. 10 µL of the cleaned PCR reaction was then circularized with KLD enzyme mix (New England Biolabs M0554S) in a 54 µL reaction. The circularized product was then cleaned with the DNA Clean & Concentrator-5 kit, eluting in 27 µL MGW.

We then transformed the cleaned assembly into commercial TransforMax EC100D pir-116 Electrocompetent *E. coli* (Biosearch Technologies EC6P095H), which has a higher transformation efficiency than our conjugative donor *E. coli* BW29427. We performed 10 individual 50 µL electroporations with 2.5 µL input DNA each, pooling all the transformations together after a 2 hour 37°C recovery. We performed liquid selection in 500 mL LB supplemented with 100 µg/mL CARB at 37°C for ∼9 hours (reached OD_600_ ∼0.6). We estimated a yield of ∼1 x 10^7^ barcodes from this transformation based on colony counts from a dilution series. We then prepared DNA using the QIAprep Spin Miniprep kit, using 9 mL input culture each for six preparations, and concentrated the pooled volume with the DNA Clean & Concentrator-5 kit, eluting in 25 µL MGW (∼77 ng/µL). The cleaned plasmid was then electroporated into *E. coli* BW29427. Six individual 50 µL electroporations with 2.5 µL input DNA were performed in total, with all transformations pooled together after a 1 hour 37°C recovery. We performed liquid selection in 375 mL LB, both supplemented with 60 µg/mL DAP and 100 µg/mL CARB at 37°C for ∼5 hours (reached OD_600_ ∼0.6). We estimated ∼4 x 10^7^ transformants in BW29427, exceeding the estimated number of barcodes. Chao1 estimates of the barcode library in the plasmid library and the BW29427 transformant library were 1.2 x 10^7^ barcodes from ∼12 million reads and 1 x 10^7^ barcodes from ∼8 million reads, respectively, corroborating the original estimates from the colony counts. The barcoded donor library was cryo-stocked in 1 mL aliquots supplemented with glycerol to a 25% (w/v) final concentration.

#### Delivery of barcoded transposons

The prepared donor library was then used to deliver the randomly barcoded transposon plasmid library to *C. testosteroni*. 864 µL (volume calculated based on OD_600_ reading before glycerol stock preparation to have ∼10X cell input coverage of the barcode diversity) of a thawed aliquot of the donor library was added to 50 mL LB, both supplemented with 60 µg/mL DAP and 100 µg/mL CARB. A 5 mL NB medium culture was started from a glycerol stock of *C. testosteroni*, and both cultures were grown at 30°C for ∼18 hours. Then, backdilutions were prepared by diluting 1 mL of the two saturated overnight cultures into 50 mL of their respective media. Both backdilutions were grown - BW29427 at 37°C and *C. testosteroni* at 30°C - for another ∼4 hours, reaching an OD_600_ of 1.3 and 0.9, respectively. The cultures were pelleted by spinning at 5,000 RCF for 10 minutes at 4°C and then washed with 1 mL PBS before being pelleted again and resuspended in PBS with the appropriate volume for a final OD_600_ of 100. Donor and recipient were mixed at a 1:1 ratio by adding 50 µL of each to an NB 1.5% agar plate (15 mm) supplemented with 60 µg/mL DAP. The conjugation was incubated for 20 hours at 30°C before being scraped with 1 mL of PBS. The scraped conjugation volume was diluted ten-fold, and 100 µL of the dilution was plated onto 32 large (245 mm) square petri dishes containing NB 1.5% agar supplemented with 200 µg/mL KAN. Plates were incubated overnight for colonies to appear. The mutant library was harvested by scraping and pooling transconjugants from selective plates in a final volume of ∼18 mL. Colonies counted using a dilution series estimated the library size to be ∼1 x 10^6^ total transconjugants. 20 µL aliquots of scraped material were saved for gDNA extraction, and the mutant library was cryo-stocked in 1 mL aliquots supplemented with glycerol to a 25% (w/v) final concentration.

### Random barcode transposon insertion sequencing

Two different library preparations were used for the random barcode Tn-seq of randomly barcoded transposon mutant library samples. The MmeI Tn-seq libraries (MmeI RB-TnSeq) can only be used with Himar1 transposon mutants, while the semi-arbitrary PCR libraries (SemiArb RB-TnSeq, **Supplementary Figure 8**) can be used with both Himar1 and Tn5 transposon mutants. gDNA extraction of transposon mutant library samples was done with the DNeasy Blood & Tissue Kit.

#### Preparation of MmeI RB-TnSeq libraries

The preparation of random barcode MmeI libraries is the same as the “Preparation of MmeI VB-TnSeq libraries” method described above, with one modification to the first PCR amplification step: the JL0003 primer (**Supplementary Table 4**) was used instead of the listed oCG0039 primer. The annealing temperature of the thermal cycling protocol was accordingly adjusted to 71°C. All other parts of the first PCR amplification and all other steps remained the same.

#### Preparation of SemiArb RB-TnSeq libraries

The preparation of random barcode semi-arbitrary PCR libraries (**Supplementary Figure 8**) is the same as the “Preparation of SemiArb VB-TnSeq libraries” method described above, with the following modifications. In the first PCR amplification step, the JL0068 primer was used instead of the listed JL0073 primer (**Supplementary Table 4**); all other parts of the first PCR amplification remained the same. The second PCR amplification step also had a primer substitution, where the JL0003 primer was used instead of the listed JL0048 primer. All other parts of the second PCR amplification and all other unmentioned steps remained the same.

#### Sequencing of RB-TnSeq libraries

DNA libraries were quantified using the Qubit™ 1X dsDNA High Sensitivity Kit and then sequenced on an Illumina NovaSeq 6000, returning a total of ∼1100 million 2 x 150 paired-end reads.

### Random barcode transposon insertion library analysis

RB-TnSeq library analysis for the characterization of the *C. testosteroni* mutant library was completed as follows. Reads were trimmed for quality and adapters using BBDuk (BBTools package) with parameters described above in the section “Variant barcode transposon insertion library analysis.” For MmeI RB-TnSeq libraries, cutadapt v5.1^43^ was used to extract genomic insertion sequences in reads with the parameters: --error-rate 0.25, -g tagaccggggacttatcatccaacctgtta, --minimum-length 10, --overlap 4. Genomic sequences were then mapped to the assembled *C. testosteroni* KF-1 genome (NCBI accession GCF_034479025) using bowtie2 v2.5.4^45^ and default parameters. A custom script using the Pysam library (see **Data and Resource Availability**) filtered reads that mapped with a MAPQ score >1 (uniquely mapped) with ≥85% identity to the reference genome. The random barcodes present in each read were extracted using Cutadapt with parameters: --error-rate 0.25, -g ATCGTCACAGGTGGAGCACTTCCG, --minimum-length 15, --overlap 4, followed by a second round of cutadapt with parameters --error-rate 0.25, -a GCGAACATACCGACCCACCCTCAT, --minimum-length 15, --maximum-length 30, --overlap 4. Barcodes were then filtered using an approach similar to Wetmore et al. 2015^46^: a barcode had to map to a unique location at least 75% of the time, the second best hit must be less than one eighth as frequent as the first hit, and had to be represented by a minimum of 3 uniquely mapped reads at that position, instead of 10. Barcodes were then linked to corresponding genes by searching for barcodes that both passed the cutoff and occurred in the middle 80% of a gene; in other words, ignoring barcodes that were inserted in the first and last 10% of all basepairs of that gene. Barcodes identified by MmeI sequencing and barcodes identified by SemiArb were combined into a single database of barcodes linked to insertion sites.

### Biodegradation fitness screen

The *C. testosteroni* randomly barcoded transposon mutant library generated above was functionally screened for the ability to grow on aromatic carbon sources.

#### Preparation of culture conditions

All bacterial cultures were prepared in M9 minimal media supplemented with three different primary carbon sources: 4-hydroxybenzoate (4HB), terephthalate, or succinate. The M9 minimal media was prepared from 2X M9 Minimal Salts (Gibco A1374401) and 1000X Trace Metals Mix (Teknova T1001) according to the manufacturer’s instructions, with only MgSO_4_ and CaCl_2_ as the additional supplements. 100 mM of carbon was spiked into the minimal media for each condition, with the specific amount of each compound varying based on their respective number of carbon atoms (14.3 mM 4HB, 12.5 mM terephthalate, and 25 mM succinate). Working stock solutions for each chemical were prepared as follows. A 0.3125 M (25X) stock solution of terephthalate was prepared by dissolving 1.970 g disodium terephthalate (TCI America T109725G) in 30 mL MGW. A 1.43 M (100X) stock solution of 4HB was prepared by dissolving 3.431 g 4-hydroxybenzoic acid sodium salt (Thermo Scientific Chemicals 391520250) in 15 mL MGW. A 0.625 M (25X) stock solution of succinate was prepared by dissolving 3.038 g succinic acid disodium salt (Thermo Scientific Chemicals 459631000) in 30 mL MGW.

#### Culture set-up and sample collection

Dilution spot plating on nutrient broth agar was done with the transposon mutant library glycerol stock to determine the viable CFU/mL in order to more precisely know the amount of inoculum for the fitness screen. One aliquot of the glycerol stock was thawed, washed twice with 1 mL no-carbon M9 minimal medium (spinning 5,000 RCF for 10 minutes at 4°C), and resuspended in the same original volume of M9 minimal medium. Spot plate counts estimated the glycerol aliquots to be stocked at around 2 x 10^6^ CFU/mL. That estimate was used to determine the amount of a freshly thawed and washed transposon mutant library glycerol aliquot used to inoculate each culture for the fitness screen, targeting at least 1 x 10^7^ CFU/mL starting concentration. After inoculation, the remaining ∼ 500 mL of the glycerol stock was pelleted (spinning 7,500 RCF for 10 minutes at room temperature) and kept frozen at −20°C as the input sample for downstream analysis. Each primary carbon source was tested in triplicate in a 4-mL volume; a no-carbon condition was included in triplicate as a negative control, and single mutant library as well as wild-type *C. testosteroni* nutrient broth cultures were included as positive growth controls for a total of 14 cultures. Cultures were set up in a 24-deepwell plate and allowed to grow shaking at 225 RPM for 48 hours at 30°C. Additional dilution spot plating on nutrient broth agar was done with the washed glycerol aliquot used to seed the cultures (done at the time of inoculation) as well as one endpoint replicate of each of the different carbon source cultures to calculate the amount of expansion across the entire growth screen for each condition: 4HB went through ∼3 doublings, terephthalate ∼11 doublings, and succinate ∼6 doublings. After spot plating, 1.5 mL of each culture was pelleted (spinning 7,500 RCF for 10 minutes at room temperature) and kept frozen at −20°C as the endpoint samples for downstream analysis.

### Random barcode sequencing

gDNA extraction of both input and endpoint samples from the biodegradation fitness screen was performed using the DNeasy Blood and Tissue Kit as described in ‘General molecular biology techniques’ above.

#### Preparation of BarSeq libraries

The first PCR amplification step was a standard KAPA HiFi HotStart PCR Kit with EvaGreen Dye qPCR reaction in a 20 µL volume according to the manufacturer’s protocol using JL0003 and JL0004 primers, 500 ng input template, and 1 µL EvaGreen dye (20X) spiked in (**Supplementary Table 4**). The reaction was completed in a qTOWER^3^G thermal cycler using a standard two-step thermal cycling protocol according to manufacturer’s instructions: 72°C annealing and extension temperature for 30 seconds. The progress of each sample was tracked through the amplification curves produced in the qPCR real-time, and the reaction was halted upon seeing sure signs of amplification above the reaction’s baseline - typically around 16-18 cycles.

Next, 1 µL of a 20-fold dilution of the first PCR product was added to the second PCR amplification as template for another standard KAPA HiFi HotStart PCR Kit with EvaGreen Dye qPCR reaction done in a 20 µL volume exactly according to the manufacturer’s protocol using i5xx and i7xx primers and 1 µL EvaGreen dye (20X). The reaction was completed in a qTOWER^3^G thermal cycler using a standard thermal cycling protocol according to the manufacturer’s instructions: 70°C annealing temperature and 15 second extension time. The progress of each sample was tracked through the amplification curves produced in the qPCR real-time, and the reaction was halted upon reaching the stage of exponential amplification - typically around an Intensity [I] value of 50000 RFU and generally around 7-9 cycles. A double-sided AMPure XP bead cleanup was performed according to the manufacturer’s protocol using an initial 0.4X bead to sample volume ratio and then a 1.2X bead to sample volume ratio to enrich for DNA fragments between 150-250 bp.

#### Sequencing of BarSeq libraries

DNA libraries were quantified using the Qubit™ 1X dsDNA High Sensitivity Kit and then sequenced on an Illumina NovaSeq 6000, returning a total of ∼1100 million 2 x 150 paired-end reads.

### BarSeq library analysis

BarSeq library analysis for the fitness screens was completed as follows. Reads were trimmed for quality and adapter using BBDuk (BBTools package) and the parameters described above. Reads were merged with BBmerge with default parameters. Barcode sequences were removed using Cutadapt v5.1^43^ with the parameters: -trimmed-only, -a CGGAAGTGCTCCACCTGTGACGAT, -g ATGAGGGTGGGTCGGTATGTTCGC, -n 2, --error-rate 0.25, --minimum-length 15, --overlap 10. Using VSEARCH v2.29^44^, barcodes were matched to the database of barcodes linked to insertions by RB-TnSeq with the parameters: --search_exact, --minseqlength 19, --maxseqlength 21. Statistical analysis of fitness results was performed using custom Python scripts. Barcode abundances for each sample were normalized to the 95% trimmed mean to account for uneven library sequencing depths and library skew. 0 values within samples were replaced with ‘pseudocounts’ of the minimum non-zero value observed across the dataset after normalization. Calculation of gene fitness was performed by taking the median abundance of each barcode across all replicates and then taking the median of all barcoded strains linked to a given gene. Fitness was calculated as log_2_ of the abundances of all strains linked to a gene within a given condition minus log_2_ of the abundances of the strains linked to a gene in the three replicates of the input libraries. A Mann-Whitney U test was performed to identify statistically significant differences in both fold change and fitness values when comparing the TER and 4HB condition to the succinate controls, and p-values were corrected for multiple hypotheses using the Benjamini-Hochberg Procedure^47^ (FDR=5%).

## Data and Resource Availability

Annotated plasmid sequences from this study are available as GenBank files under ‘Supplementary material’. All transposon/ABR promoter part plasmids and transposon plasmid destination vectors used in this study have been made available via Addgene (see **Supplementary Data Table 1** and **Supplementary Data Table 2** for Addgene IDs). In addition we are in the process of making all plasmids available as a single Addgene toolkit “Cultivarium JERBOA Toolkit”. Code for processing transposon sequencing data is made available at github.com/cultivarium/Transposon-variant-screen/.

## Author Contributions

N.O. and H.H.L. conceived of and designed the project. A.C.C. and T.P.B. performed promoter variant selection. A.C.C. and C.G. designed plasmid constructs. C.G., S.C.K. and J.L. assembled plasmid libraries and constructs. C.G. and C.K. performed transposon libraries validation experiments. S.L.B. and M.N. developed the automated, high-throughput conjugation screen. A.E., K.F. and Z.M.M. performed transposon library screening experiments. J.L. and A.C.C. developed Semi-Arb Tn-seq methods. J.L. and S.C.K. performed all NGS workflows. A.C.C. and K.M. performed all Tn-seq/NGS data analyses. J.L. generated randomly-barcoded *C. testosteroni* genome-wide mutant libraries and performed growth coupled fitness Tn-seq experiments. S.L.B. and all other authors assisted with writing the manuscript. All authors have read and approved the manuscript.

## Supporting information

Plasmid sequences

## Acknowledgements

We thank all members of the Cultivarium team for discussions throughout this project. Cultivarium acknowledges support from Eric and Wendy Schmidt as a Convergent Research Focused Research Organization (FRO).

## bioRxiv usage

Cultivarium manuscripts are disseminated as bioRxiv preprints. Comments and feedback are encouraged in the sections provided by bioRxiv.

