## Supplementary Materials for "A scalable transposon mutagenesis system for non-model bacteria"

**Supplementary Tables** **2**

Supplementary Table 1. Antibiotic resistance marker genes used in this study. 2

Supplementary Table 2. 92 bacterial strains tested using transposon variant screen (96-well plate, low-input). 3

Supplementary Table 3. Details of key plasmids and libraries used in this work. 5

Supplementary Table 4. Primers used in this study. 7

**Supplementary Figures** **8**

Supplementary Figure 1. Depiction of the transposon-insertion site sequencing approach used for transposon variant library screening. 8

Supplementary Figure 2. Overview of the transposon variant library cloning process. 9

Supplementary Figure 3. Abundance of individual variants with the Himar1-only transposon variant libraries. 10

Supplementary Figure 4. Validation of Himar1 transposon variant library performance in *E. coli* DH10B and *Pseudomonas alcaliphila*. 11

Supplementary Figure 5. Depiction of the automated 96-well plate format bacterial conjugation process used in the low-input transposon variant screen. 12

Supplementary Figure 6. Fold increase in relative promoter abundances of the top-performing promoter compared to the default endogenous promoter for each strain. 13

Supplementary Figure 7. Abundance of individual variants within the dual-transposon variant libraries. 14

Supplementary Figure 8. Depiction of the SemiArb PCR library preparation method. 15

Supplementary Figure 9. Replicability of the dual-transposase variant screen. 16

Supplementary Figure 10. *Comamonas testosteroni* transposase and KAN selectable marker promoter efficiencies. 17

Supplementary Figure 11. Genome-wide bias in insertion site frequency. 18

**Supplementary Data Files** **19**

Supplementary Data Table 1. Promoter variants used in transposon variant libraries. 19

Supplementary Data Table 2. Details of transposon destination vector plasmids used in transposon variant libraries. 19

Supplementary Data Table 3. Details of the composition and construction of all transposon variant libraries described in this study. 19

Supplementary Data Table 4. Details of the composition and construction of individual transposon variant plasmids described in this study. 19

Supplementary Data Table 5. Strains, media, and antibiotics screened in this study. 19

Supplementary Data Table 6. Promoter efficiencies and insertion counts for all samples sequenced by Tn-seq. 19

Supplementary Data Table 7. Abundance, fitness, and significance values for all *C. testosteroni* genes during the terephalate growth experiment. 19

Supplementary Data Table 8. Abundance, fitness, and significance values for all *C. testosteroni* genes during the 4-hydroxybenzoate growth experiment. 19

**References** **20**

### Supplementary Tables

##### Supplementary Table 1. Antibiotic resistance marker genes used in this study.

| Gene name | Abbreviation | Antibiotic | Source organism | Reference |
|---|---|---|---|---|
| *aph(3')-I* | KAN | Kanamycin | *Escherichia coli* | [1] |
| *aac(3)-Ia* | GEN | Gentamicin | *Pseudomonas aeruginosa* | [2] |
| *ermC* | ERM | Erythromycin | *Staphylococcus aureus* | [3] |
| *tetL* | TET | Tetracycline | *Bacillus cereus* | [3] |
| *catP* | CAM | Chloramphenicol | *Clostridium perfringens* | [4] |
| *aad(9)* | SPEC | Spectinomycin | *Enterococcus faecalis* | [4] |

**Supplementary Table 2. 92 bacterial strains tested using transposon variant screen (96-well plate, low-input).**

| Strain | Source | Taxonomy (Phylum_Class) | Max normalized OD under selection | Max Tn-seq insertion counts |
|---|---|---|---|---|
| *Cupriavidus necator* | ATCC 17699 | Pseudomonadota_Gammaproteobacteria | 0.95 | 7381 |
| *Comamonas testosteroni* | DSMZ 14576 | Pseudomonadota_Gammaproteobacteria | 0.77 | 5694 |
| *Herbaspirillum frisingense* | DSMZ 13128 | Pseudomonadota_Gammaproteobacteria | 1.03 | 4067 |
| *Shewanella putrefaciens* | ATCC 51753 | Pseudomonadota_Gammaproteobacteria | 1.04 | 1374 |
| *Methylorubrum extorquens* | ATCC 43645 | Pseudomonadota_Alphaproteobacteria | 0.69 | 893 |
| *Verrucomicrobium spinosum* | DSMZ 4136 | Verrucomicrobiota_Verrucomicrobiae | 0.62 | 693 |
| *Ectopseudomonas alcaliphila* | ATCC BAA-571 | Pseudomonadota_Gammaproteobacteria | 1.37 | 519 |
| *Sinorhizobium meliloti* | ATCC 9930 | Pseudomonadota_Alphaproteobacteria | 1.19 | 502 |
| *Shewanella oneidensis* | ATCC 700550 | Pseudomonadota_Gammaproteobacteria | 1.08 | 308 |
| *Agrobacterium fabrum* | ATCC BAA-101 | Pseudomonadota_Alphaproteobacteria | 0.84 | 230 |
| *Sphingomonas desiccabilis* | ATCC BAA-1041 | Pseudomonadota_Alphaproteobacteria | 1.74 | 173 |
| *Agrobacterium tumefaciens* | ATCC 15955 | Pseudomonadota_Alphaproteobacteria | 1.07 | 163 |
| *Anaerobutyricum hallii* | ATCC 27751 | Bacillota_A_Clostridia | 1.19 | 119 |
| *Duganella zoogloeoides* | ATCC 25935 | Pseudomonadota_Gammaproteobacteria | 2.12 | 74 |
| *Cupriavidus necator* | ATCC 43291 | Pseudomonadota_Gammaproteobacteria | 1.42 | 73 |
| *Salinivibrio costicola* | ATCC baa-952 | Pseudomonadota_Gammaproteobacteria | 1.65 | 52 |
| *Azospirillum brasilense* | ATCC 29145 | Pseudomonadota_Alphaproteobacteria | 0.77 | 51 |
| *Sphingomonas elodea* | ATCC 31461 | Pseudomonadota_Alphaproteobacteria | 0.67 | 48 |
| *Azotobacter chroococcum* | ATCC 9043 | Pseudomonadota_Gammaproteobacteria | - | 45 |
| *Shewanella electrodiphila* | ATCC baa-2408 | Pseudomonadota_Gammaproteobacteria | 1.09 | 42 |
| *Halomonas elongata* | ATCC 33173 | Pseudomonadota_Gammaproteobacteria | 1.47 | 35 |
| *Halalkalibacterium halodurans* | ATCC 21591 | Bacillota_Bacilli | 1.25 | 35 |
| *Pseudomonas synxantha* | ATCC 9890 | Pseudomonadota_Gammaproteobacteria | 1.01 | 33 |
| *Sporosarcina pasteurii* | ATCC 11859 | Bacillota_Bacilli | 0.81 | 32 |
| *Kangiella aquimarina* | DSMZ 16071 | Pseudomonadota_Gammaproteobacteria | 0.35 | 32 |
| *Rhodococcus opacus* | DSMZ 43205 | Actinomycetota_Actinomycetes | 0.3 | 27 |
| *Eubacterium limosum* | DSMZ 107592 | Bacillota_A_Clostridia | 1.21 | 25 |
| *Vibrio natriegens* | ATCC 14048 | Pseudomonadota_Gammaproteobacteria | 0.97 | 24 |
| *Chromohalobacter canadensis* | ATCC 43984 | Pseudomonadota_Gammaproteobacteria | 0.81 | 24 |
| *Streptomyces lividans* | ATCC 19844 | Actinomycetota_Actinomycetia | 0.53 | 24 |
| *Streptomyces venezuelae* | ATCC 10712 | Actinomycetota_Actinomycetia | 0.73 | 23 |
| *Chitinophaga sancti* | ATCC 23090 | Bacteroidota_Bacteroidia | 0.39 | 20 |
| *Priestia megaterium* | ATCC 14581 | Bacillota_Bacilli | 0.42 | 19 |
| *Shewanella amazonensis* | ATCC 700329 | Pseudomonadota_Gammaproteobacteria | 2.01 | 18 |
| *Clostridium sporogenes* | ATCC 3584 | Bacillota_A_Clostridia | 1.77 | 18 |
| *Idiomarina loihiensis* | ATCC baa-735 | Pseudomonadota_Gammaproteobacteria | 0.54 | 17 |
| *Zymomonas mobilis* | DSMZ 22645 | Pseudomonadota_Alphaproteobacteria | 0.3 | 16 |
| *Azospirillum lipoferum* | ATCC 29707 | Pseudomonadota_Alphaproteobacteria | 2.08 | 14 |
| *Bogoriella caseilytica* | ATCC 700413 | Actinomycetota_Actinomycetes | 2 | 14 |
| *Exiguobacterium aurantiacum* | ATCC baa-333 | Bacillota_Bacilli | 0.96 | 14 |
| *Arcobacter defluvii* | DSMZ 25359 | Campylobacterota_Campylobacteria | 0.64 | 13 |
| *Vreelandella subglaciescola* | ATCC 43669 | Pseudomonadota_Gammaproteobacteria | 0.52 | 12 |
| *Lacicoccus alkaliphilus* | ATCC baa-722 | Bacillota_Bacilli | 0.51 | 11 |
| *Ammoniphilus oxalivorans* | ATCC 700648 | Bacillota_Bacilli | 0.85 | 10 |
| *Streptomyces albidoflavus* | ATCC 23899 | Actinomycetota_Actinomycetes | 2.52 | 8 |
| *Vreelandella neptunia* | ATCC BAA-805 | Pseudomonadota_Gammaproteobacteria | 1.94 | 8 |

| Strain | Source | Taxonomy (Phylum_Class) | Max normalized OD under selection | Max Tn-seq insertion counts |
|---|---|---|---|---|
| *Azotobacter vinelandii* | ATCC 478 | Pseudomonadota_Gammaproteobacteria | 1.21 | 7 |
| *Lactococcus lactis* | ATCC 19435 | Bacillota_Bacilli | 0.42 | 6 |
| *Cellulophaga lytica* | DSMZ 7489 | Bacteroidota_Bacteroidia | 3.08 | 4 |
| *Marinococcus halophilus* | ATCC 27964 | Bacillota_Bacilli | 0.34 | 3 |
| *Blautia wexlerae* | ATCC BAA-1564 | Bacillota_A_Clostridia | 0.7 | 2 |
| *Rhodococcus erythropolis* | ATCC 4277 | Actinomycetota_Actinomycetes | 0.58 | 1 |
| *Paenibacillus polymyxa* | ATCC 842 | Bacillota_Bacilli | 0.51 | 1 |
| *Lysinibacillus fusiformis* | ATCC 7055 | Bacillota_Bacilli | 0.8 | 0 |
| *Evansella cellulosilytica* | ATCC 21833 | Bacillota_Bacilli | 0.63 | 0 |
| *Deinococcus radiodurans* | ATCC 13939 | Deinococcota_Deinococci | 0.62 | 0 |
| *Azospirillum lipoferum* | ATCC 29707 | Pseudomonadota_Alphaproteobacteria | 2.08 | - |
| *Piscinibacter sakaiensis* | DSMZ 112585 | Pseudomonadota_Gammaproteobacteria | 1.23 | - |
| *Agrobacterium tumefaciens* | ATCC 15955 | Pseudomonadota_Alphaproteobacteria | 1.07 | - |
| *Photorhabdus luminescens* | ATCC 29999 | Pseudomonadota_Gammaproteobacteria | 0.92 | - |
| *Rothia mucilaginosa* | ATCC 25296 | Actinomycetota_Actinomycetia | 0.9 | - |
| *Agrobacterium fabrum* | ATCC BAA-101 | Pseudomonadota_Alphaproteobacteria | 0.84 | - |
| *Methylobacterium radiotolerans* | ATCC 27329 | Pseudomonadota_Alphaproteobacteria | 0.82 | - |
| *Lacrimispora saccharolytica* | ATCC 35040 | Bacillota_A_Clostridia | 0.81 | - |
| *Flavobacterium johnsoniae* | ATCC 17061 | Bacteroidota_Bacteroidia | 0.7 | - |
| *Intestinibacter bartlettii* | DSMZ 16795 | Bacillota_A_Clostridia | 0.7 | - |
| *Alkalihalophilus pseudofirmus* | ATCC 700159 | Bacillota_Bacilli | 0.61 | - |
| *Enterocloster bolteae* | ATCC BAA-613 | Bacillota_A_Clostridia | 0.61 | - |
| *Gemmatimonas aurantiaca* | DSMZ 14586 | Gemmatimonadota_Gemmatimonadetes | 0.61 | - |
| *Azospirillum halopraeferens* | DSMZ 3675 | Pseudomonadota_Alphaproteobacteria | 0.58 | - |
| *Agathobacter rectalis* | ATCC 33656 | Bacillota_A_Clostridia | 0.45 | - |
| *Shewanella indica* | ATCC BAA-2732 | Pseudomonadota_Gammaproteobacteria | 0.44 | - |
| *Eubacterium ramulus* | ATCC 29099 | Bacillota_Clostridia | 0.37 | - |
| *Streptomyces violaceoruber* | ATCC BAA-471 | Actinomycetota_Actinomycetia | 0.36 | - |
| *Alistipes shahii* | ATCC BAA-1179 | Bacteroidota_Bacteroidia | - | - |
| *Alkalibacterium sp. TRTYP6* | ATCC baa-2622 | Bacillota_Bacilli | - | - |
| *Armatimonas rosea* | DSMZ 23562 | Armatimonadota_Armatimonadia | - | - |
| *Bacillus licheniformis* | ATCC 14580 | Bacillota_Bacilli | - | - |
| *Eubacterium limosum* | ATCC 8486 | Bacillota_A_Clostridia | - | - |
| *Komagataeibacter rhaeticus* | ATCC BAA-2831 | Pseudomonadota_Alphaproteobacteria | - | - |
| *Komagataeibacter sucrofermentans* | ATCC 700178 | Pseudomonadota_Alphaproteobacteria | - | - |
| *Lysinibacillus sphaericus* | ATCC 14577 | Bacillota_Bacilli | - | - |
| *Mycolicibacterium smegmatis* | ATCC 607 | Actinomycetota_Actinomycetia | - | - |
| *Oerskovia enterophila* | ATCC 35307 | Actinomycetota_Actinomycetes | - | - |
| *Planctopirus limnophila* | DSMZ 3776 | Planctomycetota;_Planctomycetia | - | - |
| *Rhizobium leguminosarum* | ATCC 10004 | Pseudomonadota_Alphaproteobacteria | - | - |
| *Ruminococcus bromii* | ATCC 27255 | Bacillota_A_Clostridia | - | - |
| *Shouchella clausii* | ATCC 700160 | Bacillota_Bacilli | - | - |
| *Sphingomonas echinoides* | ATCC 14820 | Pseudomonadota_Alphaproteobacteria | - | - |
| *Sporosarcina pasteurii* | DSMZ 33 | Bacillota_Bacilli | - | - |
| *Streptomyces albus* | ATCC 3004 | Actinomycetota_Actinomycetia | - | - |
| *Streptomyces virginiae* | ATCC 12630 | Actinomycetota_Actinomycetes | - | - |
| *Terriglobus roseus* | DSMZ 18391 | Acidobacteriota_Terriglobia | - | - |
| *Vreelandella alkaliphila* | ATCC baa-953 | Pseudomonadota_Gammaproteobacteria | - | - |
| *Xanthomonas campestris* | ATCC 33913 | Pseudomonadota_Gammaproteobacteria | - | - |

#### Supplementary Table 3. Details of key plasmids and libraries used in this work.

| Name | Description | Experiment | Transposase | ABR | Source |
|---|---|---|---|---|---|
| Single Plasmid | | | | | |
| pGL2_175 | Control plasmid for GEN transposon variant library screening | • Tn screen on large cohort of bacterial strains | None | GEN | Gilbert et al. 2024[5] |
| pGL2_228 | Control plasmid for KAN transposon variant library screening | • Tn screen on large cohort of bacterial strains | None | KAN | Gilbert et al. 2024[5] |
| pGL2_229 | Control plasmid for ERM transposon variant library screening | • Tn screen on large cohort of bacterial strains | None | ERM | Gilbert et al. 2024[5] |
| pGL2_230 | Control plasmid for TET transposon variant library screening | • Tn screen on large cohort of bacterial strains | None | TET | Gilbert et al. 2024[5] |
| pGL2_231 | Control plasmid for CAM transposon variant library screening | • Tn screen on large cohort of bacterial strains | None | CAM | Gilbert et al. 2024[5] |
| pGL2_232 | Control plasmid for SPEC transposon variant library screening | • Tn screen on large cohort of bacterial strains | None | SPEC | Gilbert et al. 2024[5] |
| pGLU_127 | Transposon variant plasmid (individually cloned) | • Best-performing transposon variant for *Comamonas testosteron*i transposon mutagenesis | Himar1 | KAN | This study |
| Plasmid Library | | | | | |
| pGLU_28 | Himar1-KAN transposon variant library | • Initial *E. coli* Tn variant validation<br>• Tn screen on large cohort of bacterial strains | Himar1 | GEN | This study |
| pGLU_29 | Himar1-GEN transposon variant library | • Initial *E. coli* Tn variant validation<br>• Tn screen on large cohort of bacterial strains | Himar1 | KAN | This study |
| pGLU_30 | Himar1-ERM transposon variant library | • Initial *E. coli* Tn variant validation<br>• Tn screen on large cohort of bacterial strains | Himar1 | ERM | This study |
| pGLU_31 | Himar1-TET transposon variant library | • Initial *E. coli* Tn variant validation<br>• Tn screen on large cohort of bacterial strains | Himar1 | TET | This study |
| pGLU_32 | Himar1-CAM transposon variant library | • Initial *E. coli* Tn variant validation<br>• Tn screen on large cohort | Himar1 | CAM | This study |

| | | | | | |
|---|---|---|---|---|---|
| | | of bacterial strains | | | |
| pGLU_33 | Himar1-SPEC transposon variant library | • Initial *E. coli* Tn variant validation<br>• Tn screen on large cohort of bacterial strains | Himar1 | SPEC | This study |
| pGLU_135 | Tn5-KAN transposon variant library | • Construction of dual-transposon variant library | Tn5 | KAN | This study |
| pGLU_136 | Tn5-GEN transposon variant library | • Construction of dual-transposon variant library | Tn5 | GEN | This study |
| pGLU_137 | Tn5-ERM transposon variant library | • Construction of dual-transposon variant library | Tn5 | ERM | This study |
| pGLU_138 | Tn5-TET transposon variant library | • Construction of dual-transposon variant library | Tn5 | TET | This study |
| pGLU_139 | Tn5-CAM transposon variant library | • Construction of dual-transposon variant library | Tn5 | CAM | This study |
| pGLU_140 | Tn5-SPEC transposon variant library | • Construction of dual-transposon variant library | Tn5 | SPEC | This study |
| pGLU_183 | Himar1/Tn5-KAN transposon variant library | • Dual-transposon variant library screening | Himar1/Tn5 | KAN | This study |
| pGLU_184 | Himar1/Tn5-GEN transposon variant library | • Dual-transposon variant library screening | Himar1/Tn5 | GEN | This study |
| pGLU_185 | Himar1/Tn5-ERM transposon variant library | • Dual-transposon variant library screening | Himar1/Tn5 | ERM | This study |
| pGLU_186 | Himar1/Tn5-TET transposon variant library | • Dual-transposon variant library screening | Himar1/Tn5 | TET | This study |
| pGLU_187 | Himar1/Tn5-CAM transposon variant library | • Dual-transposon variant library screening | Himar1/Tn5 | CAM | This study |
| pGLU_188 | Himar1/Tn5-SPEC transposon variant library | • Dual-transposon variant library screening | Himar1/Tn5 | SPEC | This study |
| pGLU_127_RB | Barcoded library for genome-wide mutagenesis | • Best-performing transposon variant for *Comamonas testosteroni* transposon mutagenesis with random barcodes for genome-wide mutant library generation | Himar1 | KAN | This study |

**Supplementary Table 4. Primers used in this study.**

Regions of homology to the fragment being amplified are shown in bold. The underlined regions represent 5’ primer extensions which introduce stub sequences enabling binding of dual-indexed Illumina i5 and i7 primers in the subsequent PCR step.

| Primer | Sequence | Description |
|---|---|---|
| pMarC9_ P5AdaptDup | 5’-ACACTCTTTCCCTACACGACGCTCTTCCGATCT(N:25252525)(N)(N)-3’<br>3’-Phos-TGTGAGAAAGGGATGTGCTGCGAGAAGGCTAGA-Phos-5’ | Oligo duplex used in MmeI VB-TnSeq and MmeI RB-TnSeq ligation step |
| i5xx | AATGATACGGCGACCACCGAGATCTACACNNNNNNNNNN**ACACTCTTTCCCTACACGACGCTCTTCCGATCT** | MmeI VB-TnSeq and RB-TnSeq PCR 1; SemiArb VB-TnSeq and RB-TnSeq PCR 2; and BarSeq PCR 2 forward primer example, used to add Illumina i5 index to the sample |
| oCG0038 | TCCCTACACGACGCTCTTCCGATCT**GGGTCACGCGTAGGACG** | VBBarSeq PCR 1 forward primer |
| oCG0039 | GAGTTCAGACGTGTGCTCTTCCGATCT**CCAGCTTCACACGGCGT** | MmeI VB-TnSeq and VBBarSeq PCR 1 reverse primer |
| P5 | **AATGATACGGCGACCACCGAGA** | MmeI VB-TnSeq and RB-TnSeq PCR 2; SemiArb VB-TnSeq and RB-TnSeq PCR 3 forward primer on the P5 Illumina adapter |
| i7xx | CAAGCAGAAGACGGCATACGAGATNNNNNNNNNNGTGACTG**GAGTTCAGACGTGTGCTCTTCCGATCT** | MmeI VB-TnSeq and RB-TnSeq PCR 2; SemiArb VB-TnSeq and RB-TnSeq PCR 3; and BarSeq PCR 2 reverse primer example, used to add Illumina i7 index to the sample |
| JL0075 | AAACACGTGGCAAACATTCC**NNNNNNNNNN** | SemiArb VB-TnSeq and RB-TnSeq primer for arbitrary priming |
| JL0076 | TCCCTACACGACGCTCTTCCGATCT**AAACACGTGGCAAACATTCC** | SemiArb VB-TnSeq and RB-TnSeq PCR 1 forward primer |
| JL0073 | **CCAGCTTCACACGGCGT** | SemiArb VB-TnSeq PCR 1 reverse primer |
| JL0048 | GAGTTCAGACGTGTGCTCTTCCGATCT**ACACGGCGTGTGGCTGCGGAAC** | SemiArb VB-TnSeq PCR 2 reverse primer |
| JL0061 | ACATACCGACCCACCCTCAT**GCGTCCTACGCGTGACCC** | Reverse primer for adding the barcodes and the orthogonal primer-binding site to pGLU_127 |
| JL0062 | TCGCNNNNNNNNNNNNNNNNNNNNCGGAAGTGCTCCACCTGTGACGAT**CACGCCGTGTGAAGCTGG** | Forward primer for adding the barcodes and the orthogonal primer-binding site to pGLU_127 |
| JL0003 | GTGACTGGAGTTCAGACGTGTGCTCTTCCGATCT**ATCGTCACAGGTGGAGCACTTCCG** | MmeI RB-TnSeq PCR 1, SemiArb RB-TnSeq PCR 2, and BarSeq PCR 1 reverse primer |
| JL0068 | **CCAGCTTCACACGGCGTGATCGTCACA** | SemiArb RB-TnSeq PCR 1 reverse primer |
| JL0004 | ACACTCTTTCCCTACACGACGCTCTTCCGATCT**ATGAGGGTGGGTCGGTATGTTCGC** | BarSeq PCR 1 forward primer |

#### Supplementary Figures

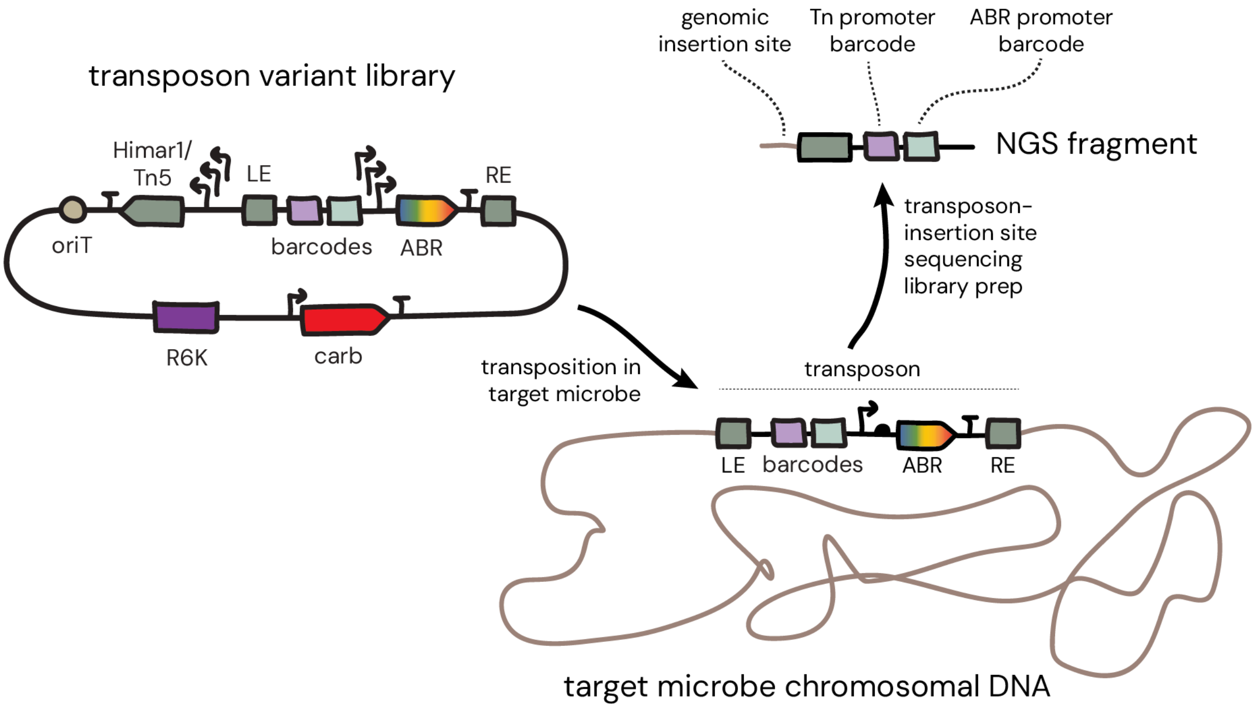


**Supplementary Figure 1. Depiction of the transposon-insertion site sequencing approach used for transposon variant library screening.**

Transposon variant libraries are delivered as a pool to the target microbe. In cases where true transposition events occur, transposons are inserted into the genome of the target microbe. Unique barcodes within the transposon carry the identity of: the promoter variant driving the antibiotic resistance (ABR) cassette (pale sage) and the promoter variant driving the transposase gene as well as the transposase enzyme variant (Himar1 or Tn5) itself (lilac). Transposon-insertion site sequencing methods (either MmeI VB-TnSeq or SemiArb VB-TnSeq) allow simultaneous capture of these barcodes and genomic insertion sites. Downstream computational pipelines use the number of observed insertion sites per unique barcode combination to calculate the relative efficiency of transposon variants. LE, left mosaic end; RE, right mosaic end; oriT, origin of transfer sequence; R6K, R6Kγ conditional origin of replication; carb, carbenicillin resistance gene marker.

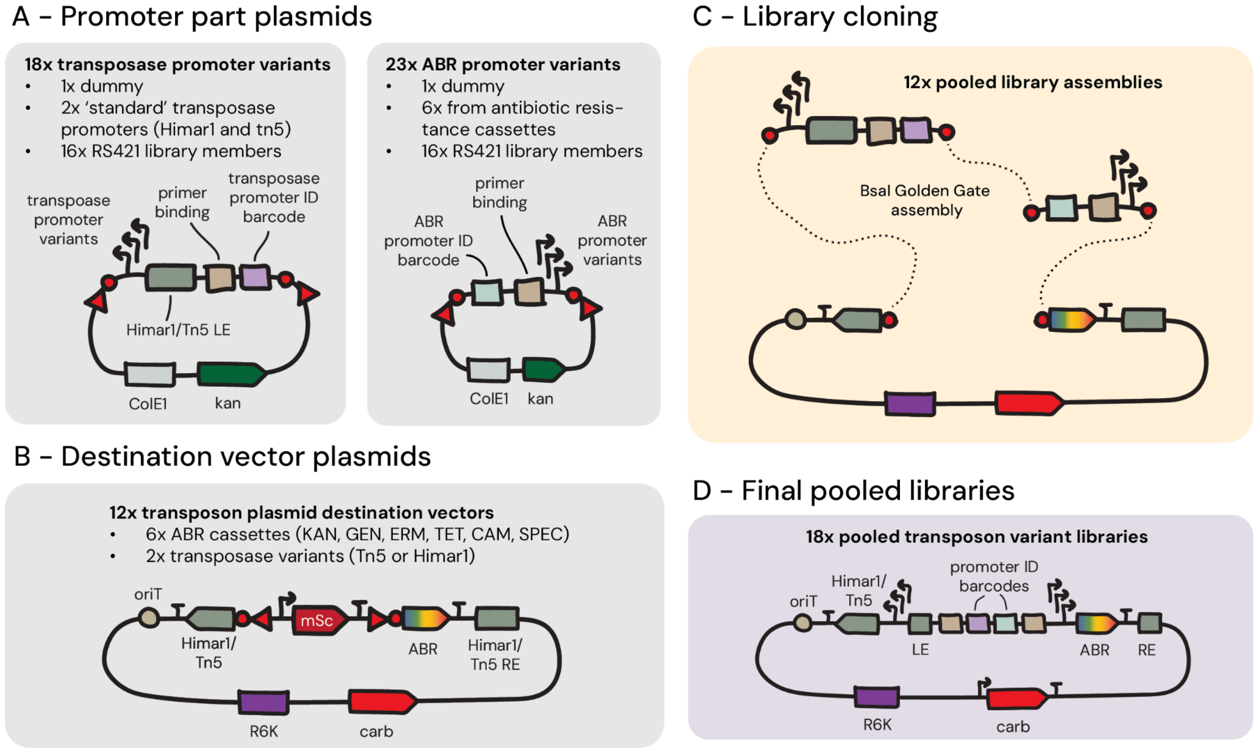


**Supplementary Figure 2. Overview of the transposon variant library cloning process.**
(**A**) Promoter part plasmid composition. All promoter variants were generated as part plasmids, where the relevant promoter region is flanked by BsaI restriction enzyme cut sites (red triangle and circle) and maintained on a standard *E. coli* plasmid for ease of propagation. 18X transposon promoter variants were generated for each transposase type (Tn5 or Himar1): each unique promoter was combined with a Himar1 or Tn5 left mosaic end (LE), a constant primer binding region, and a variable 20 bp promoter ID barcode. 23X antibiotic resistance (ABR) marker promoter variants were generated: each unique promoter was combined with a constant primer binding region and a variable 20 bp promoter ID barcode. (**B**) Transposon destination vector plasmid composition. A set of 12X unique transposon destination vector plasmids were created from all combinations of 6X possible ABR markers and 2X possible transposase enzyme variants. All plasmids possessed an RK2/RP4 origin of transfer sequence (oriT), an mScarlet reporter gene (mSc) flanked by BsaI restriction enzyme cut sites (red triangle and circle), a Himar1 or Tn5 right mosaic end (RE) sequence, the conditional *E. coli* origin of replication R6Kγ, and an *E. coli* antibiotic resistance marker (carb). (**C**) Tn-variant libraries were generated by pooling the relevant promoter part plasmids and performing BsaI Golden Gate assembly reactions into the transposon destination vector plasmids. (**D**) The general architecture of final pooled Tn-variant libraries. We built 18X final libraries: 6X Himar1-only libraries, 6X Tn5-only libraries, and 6X Himar+Tn5 libraries.

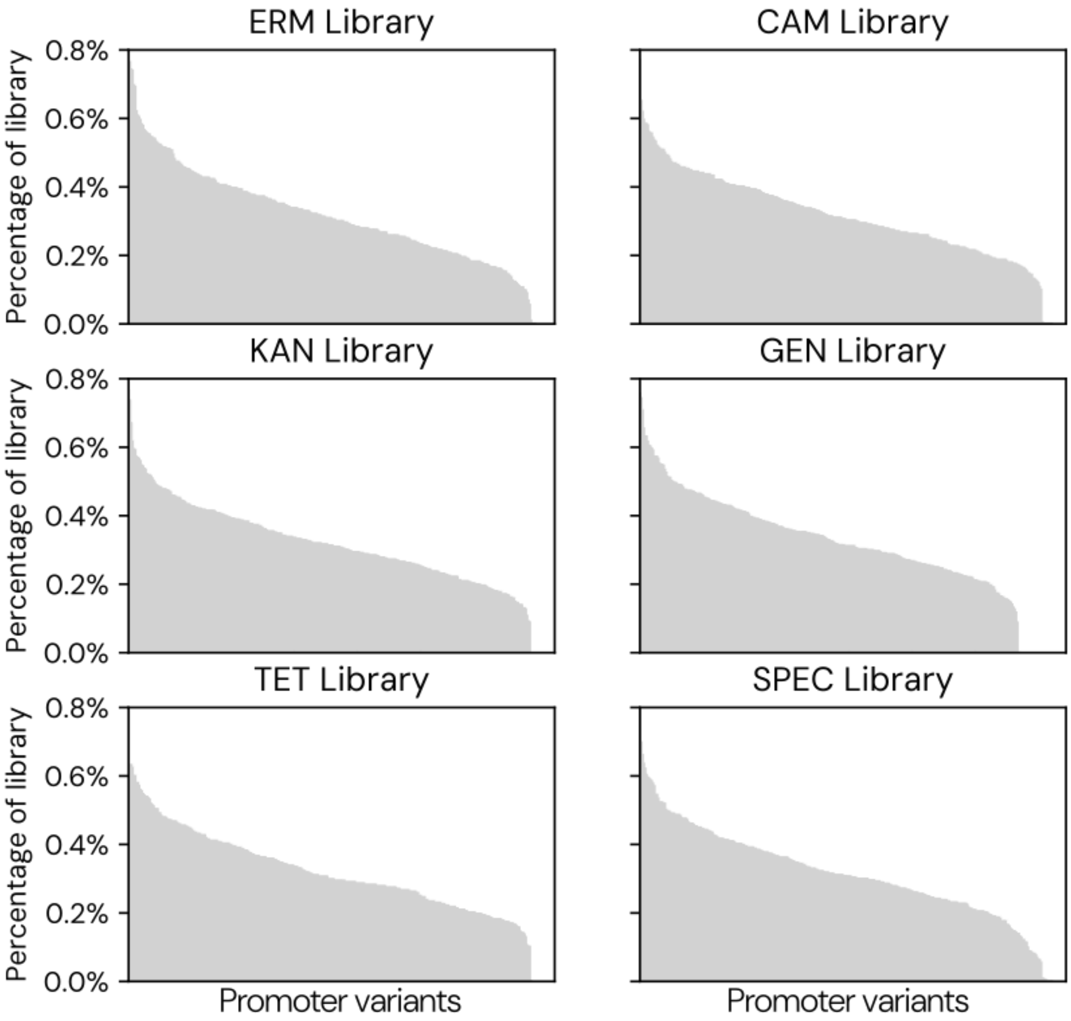


**Supplementary Figure 3. Abundance of individual variants with the Himar1-only transposon variant libraries.**

Abundances of all possible 324 combinatorial promoter variants in each of the six Himar1 transposon variant libraries, as measured by amplicon sequencing.

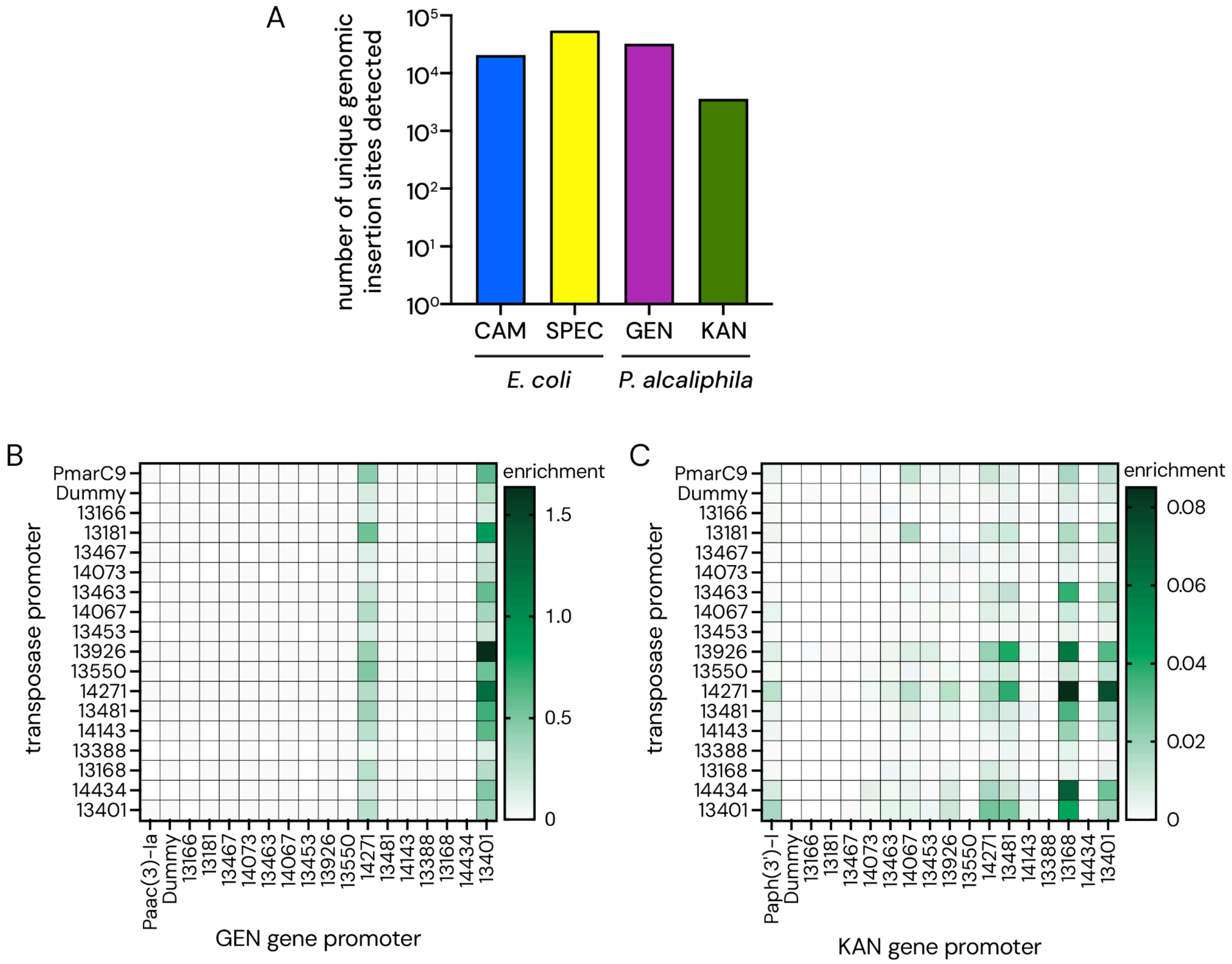


**Supplementary Figure 4. Validation of Himar1 transposon variant library performance in *E. coli* DH10B and *Pseudomonas alcaliphila*.**

(**A**) Quantification of the number of unique insertion sites observed per strain-library combination by Tn-seq. Two Himar1 transposon variant libraries were delivered to each strain: CAM and SPEC to *E. coli*, and GEN and KAN to *P. alcaliphila*. Transconjugants were pooled and analyzed by MmeI VB-TnSeq to quantify the total number of genomic insertion sites detected across all library variants. (**B-C**) Heatmaps showing the relative transposition efficiency (enrichment) of promoter variants driving expression of Himar1 and the ABR gene. Data shown from the (**B**) GEN and (**C**) KAN transposon variant libraries delivered to *P. alcaliphila*. The equivalent data for *E. coli* is shown in **Figure 1C-D**. Enrichment represents per variant unique insertion site abundance from transconjugants normalized to per variant abundance from the library hosted in the *E. coli* conjugative donor.

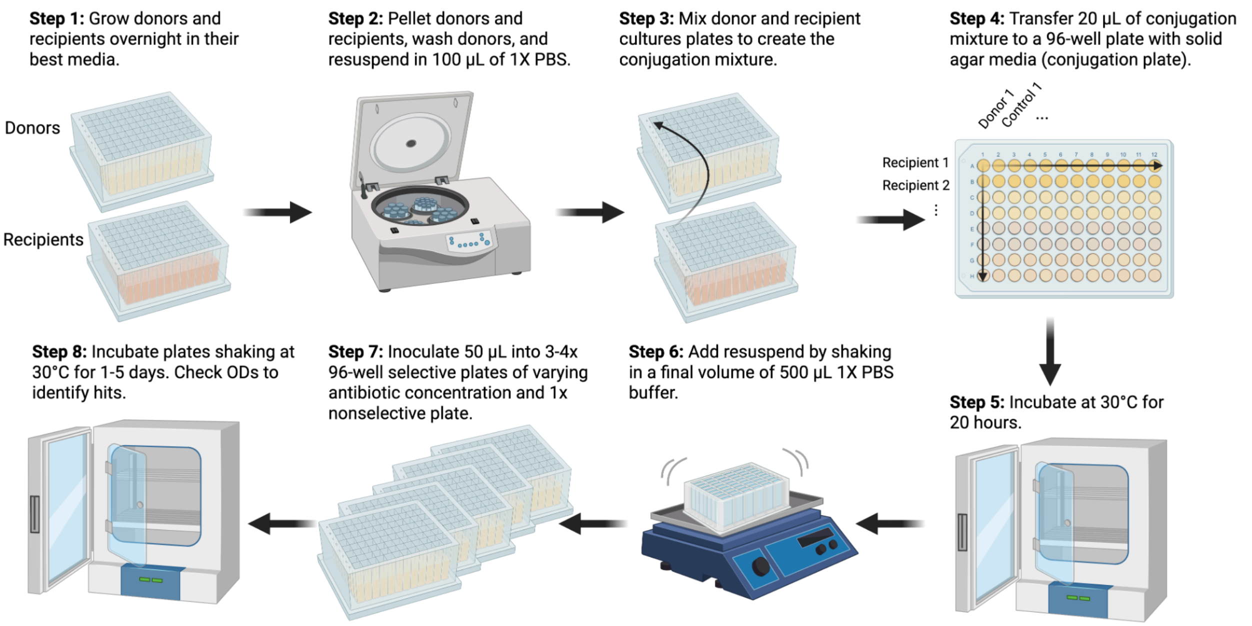


**Supplementary Figure 5. Depiction of the automated 96-well plate format bacterial conjugation process used in the low-input transposon variant screen.**

This figure illustrates the general steps performed as part of the standard 96-well plate format bacterial conjugations performed in this study. Importantly, for particular microbes, parameters such as growth temperature and time were modified as described in Methods. Step 1: transposon variant libraries cloned into *E. coli* conjugative donor strain (donors) and the bacterial target strains (recipients) are grown to generate biomass for conjugations. Step 2: donor and recipient cultures are concentrated. Donor cultures are additionally washed in PBS to remove residual medium and antibiotics. Step 3: Donor and recipient samples are combined to create the conjugation mixture. Step 4: The conjugation mixture is transferred to solid agar media in a 96-well conjugation plate. Recipient cultures are arranged by row (eight recipients per plate), and donor cultures are arranged by column (twelve donors per plate). Since one negative control donor is used for each library type, the result is a total of six possible transposon variant libraries screened against eight possible recipient strains. Step 5: co-cultures are incubated at 30°C for 20 hours to allow conjugation to occur. Step 6: biomass is resuspended from the conjugation plate wells by adding PBS and shaking. Step 7: biomass is inoculated into multiple 96-well deep well plate liquid cultures, matching the footprint of the conjugation plate: a single plate with no antibiotic and the recipient strain's preferred medium (non-selective) and three or four plates with variable concentrations of the relevant antibiotics for selection of transposon insertions. Step 8: liquid cultures are monitored over several days by $OD_{600}$ measurement; all samples for which increased growth in the donor library condition versus the control condition is observed are harvested for genomic DNA extraction and NGS analysis. Created in BioRender. Gilbert, C. (2025) https://BioRender.com/3wbdboq

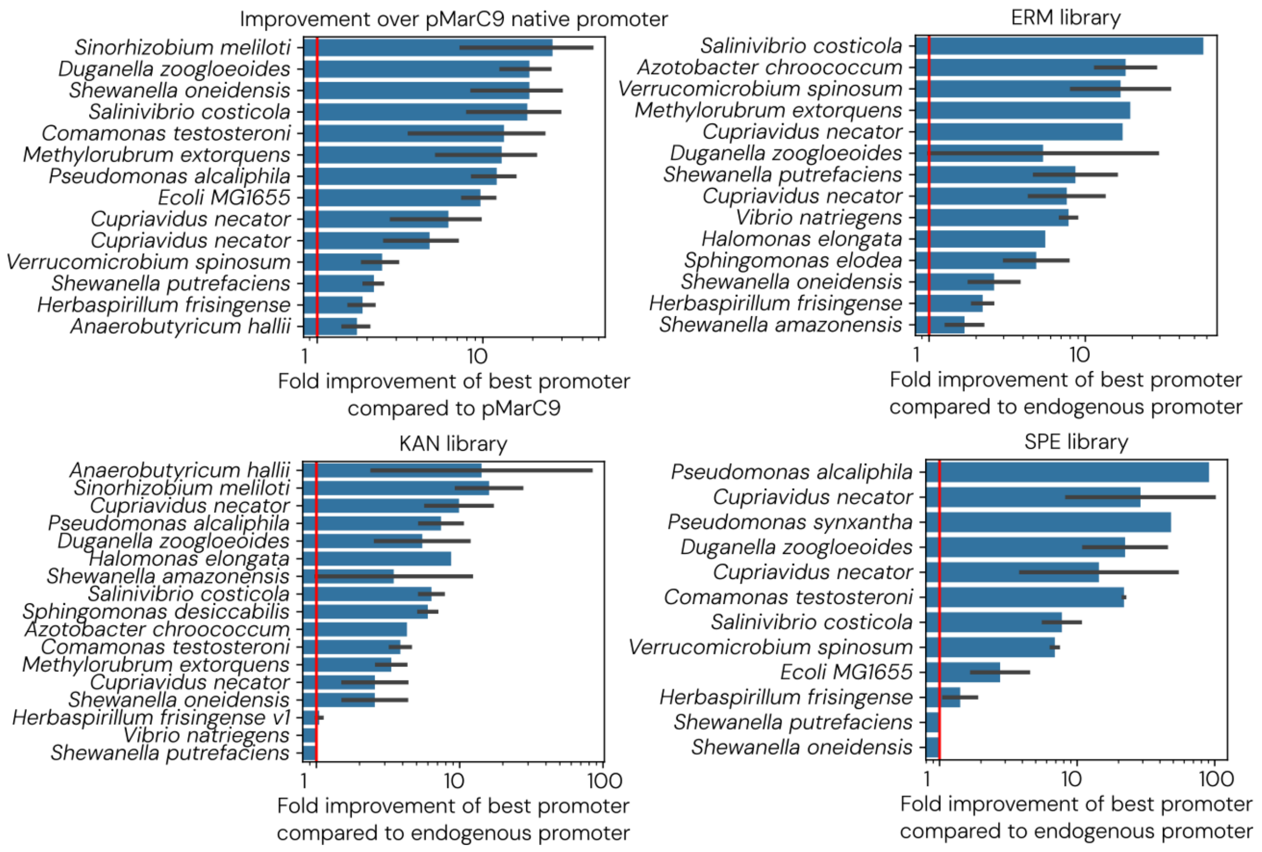


**Supplementary Figure 6. Fold increase in relative promoter abundances of the top-performing promoter compared to the default endogenous promoter for each strain.** Starting top left and moving clockwise: fold improvement of the best-performing transposase promoter variant compared to the endogenous pMarC9 transposon promoter; fold improvement of the best-performing *ermC* (ERM) gene promoter compared to the endogenous *ermC* gene promoter; fold improvement of the best-performing *aad(9)* (SPEC) gene promoter compared to the endogenous *aad(9)* gene promoter; fold improvement of the best-performing *aph(3')-I* (KAN) gene promoter compared to the endogenous *aph(3')-I* gene promoter. Red lines represent 1X - *i.e.,* identical efficiency compared to the endogenous promoters.

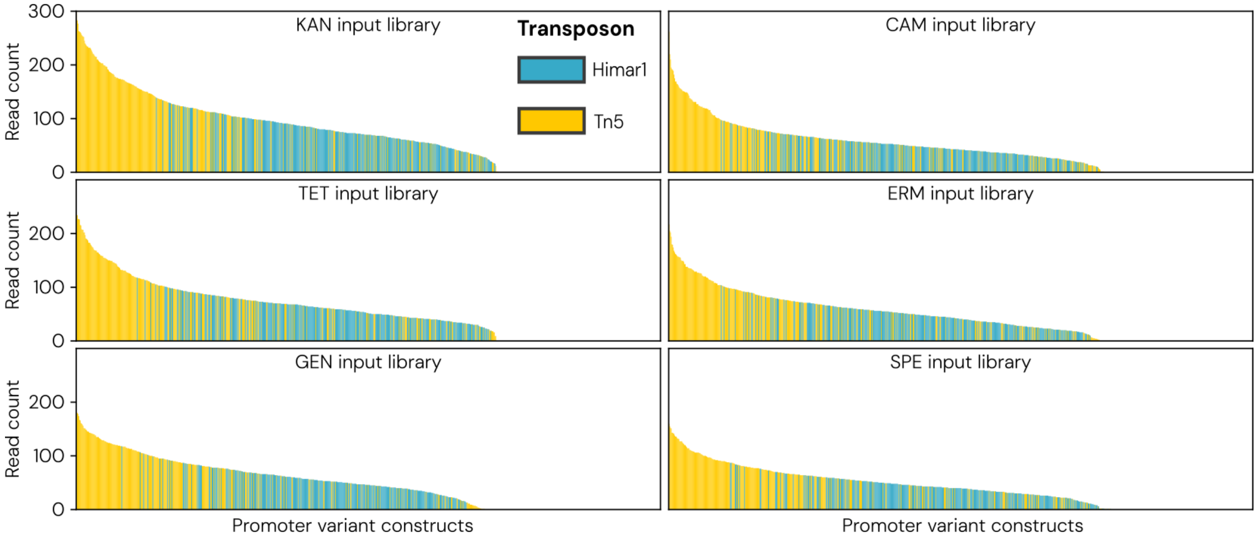


**Supplementary Figure 7. Abundance of individual variants within the dual-transposon variant libraries.**

Each bar represents the abundance of an individual member of the transposon variant libraries. Plasmid DNA was extracted from pooled libraries in the *E. coli* conjugative donor strain and sequenced by amplicon-sequencing. Himar1 variants are shown in blue and Tn5 variants are shown in yellow.

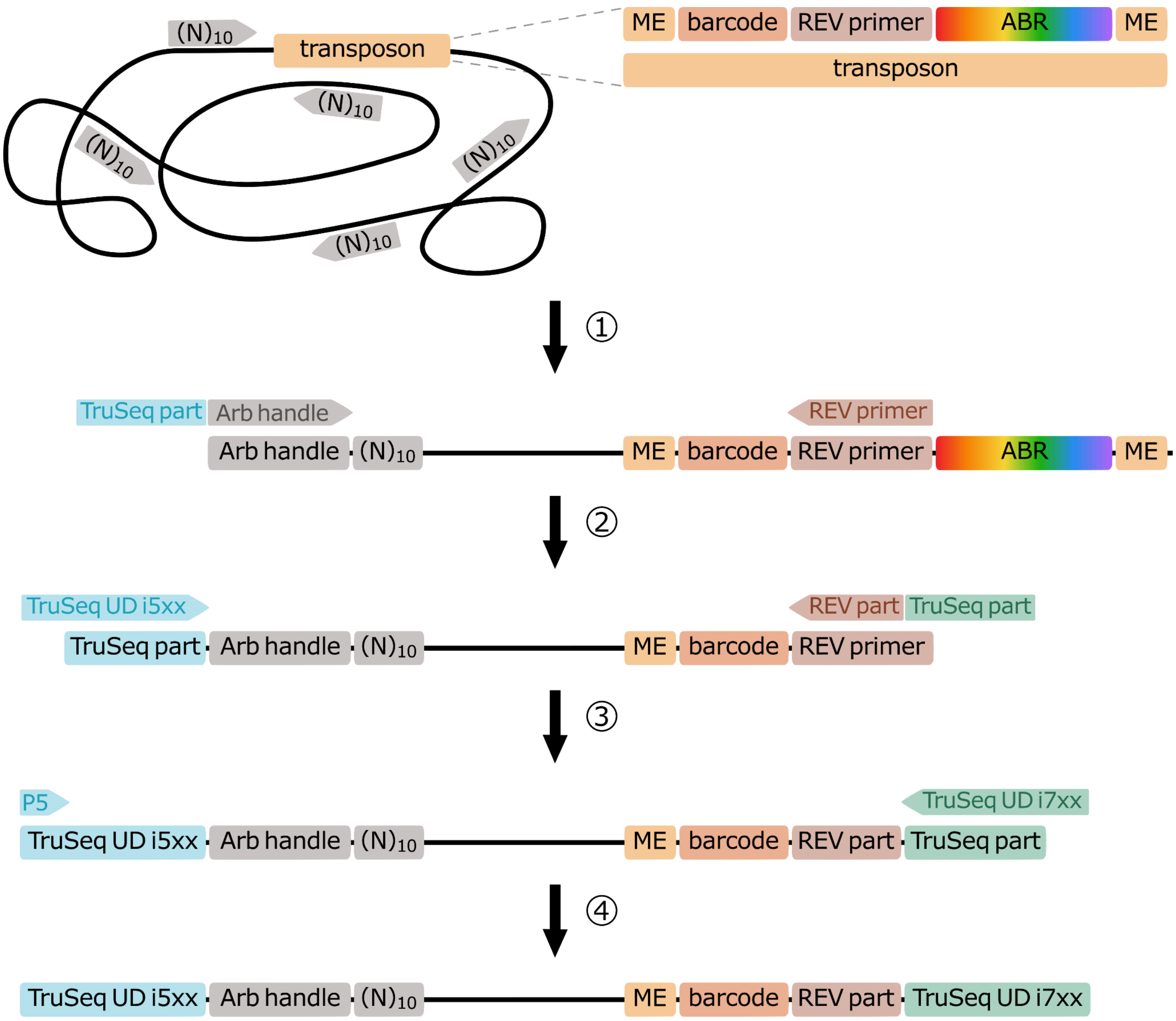


**Supplementary Figure 8. Depiction of the SemiArb PCR library preparation method.** Semi-arbitrary PCR insertion library construction process. The input is genomic DNA with a transposon containing a barcode, a specific reverse (REV) primer binding site, and an antibiotic resistance cassette (ABR) between the two mosaic ends (ME). 1) Arbitrary priming is performed through five cycles of amplification with $N_{10}$-mers oligonucleotides bearing a 5' constant sequence ("Arb handle"). 2) PCR 1 amplifies the handle of the arbitrary primer along with the conserved reverse primer binding site in the transposon itself. 3) PCR 2 continues to enrich transposon-containing arbitrary amplification products while adding the 5' adapter sequences needed for Illumina sequencing. 4) PCR 3 completes the library construction by adding the 3' Illumina adapter sequences. Lengths of the library construct are controlled by limited extension time in the arbitrary priming process, as well as size selection through bead cleaning.

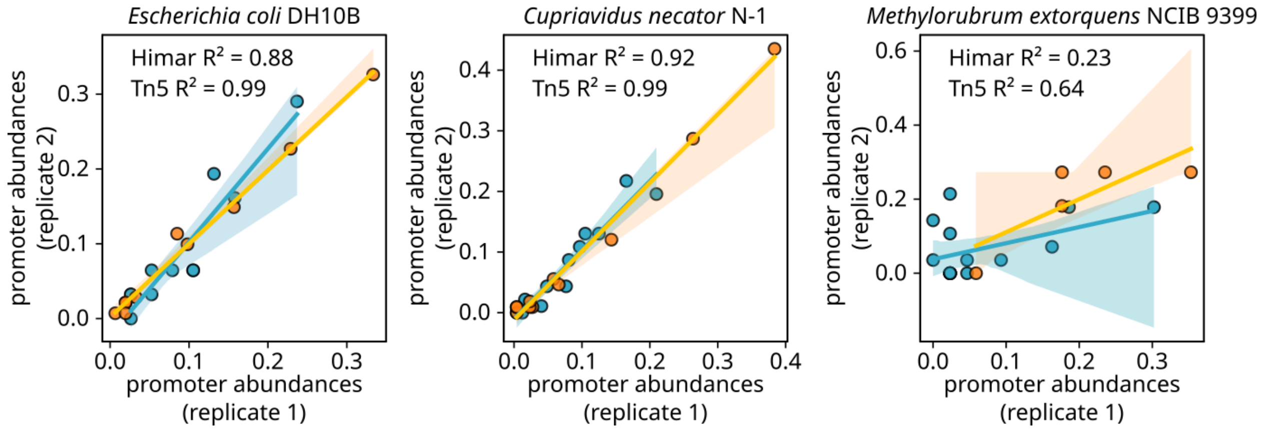


**Supplementary Figure 9. Replicability of the dual-transposase variant screen.**

For each organism, promoter abundances are compared between two biological replicates (replicate 1 shown on the x-axis, replicate 2 shown on the y-axis). Promoter abundances are shown for both Tn5 (orange) and Himar1 (blue).

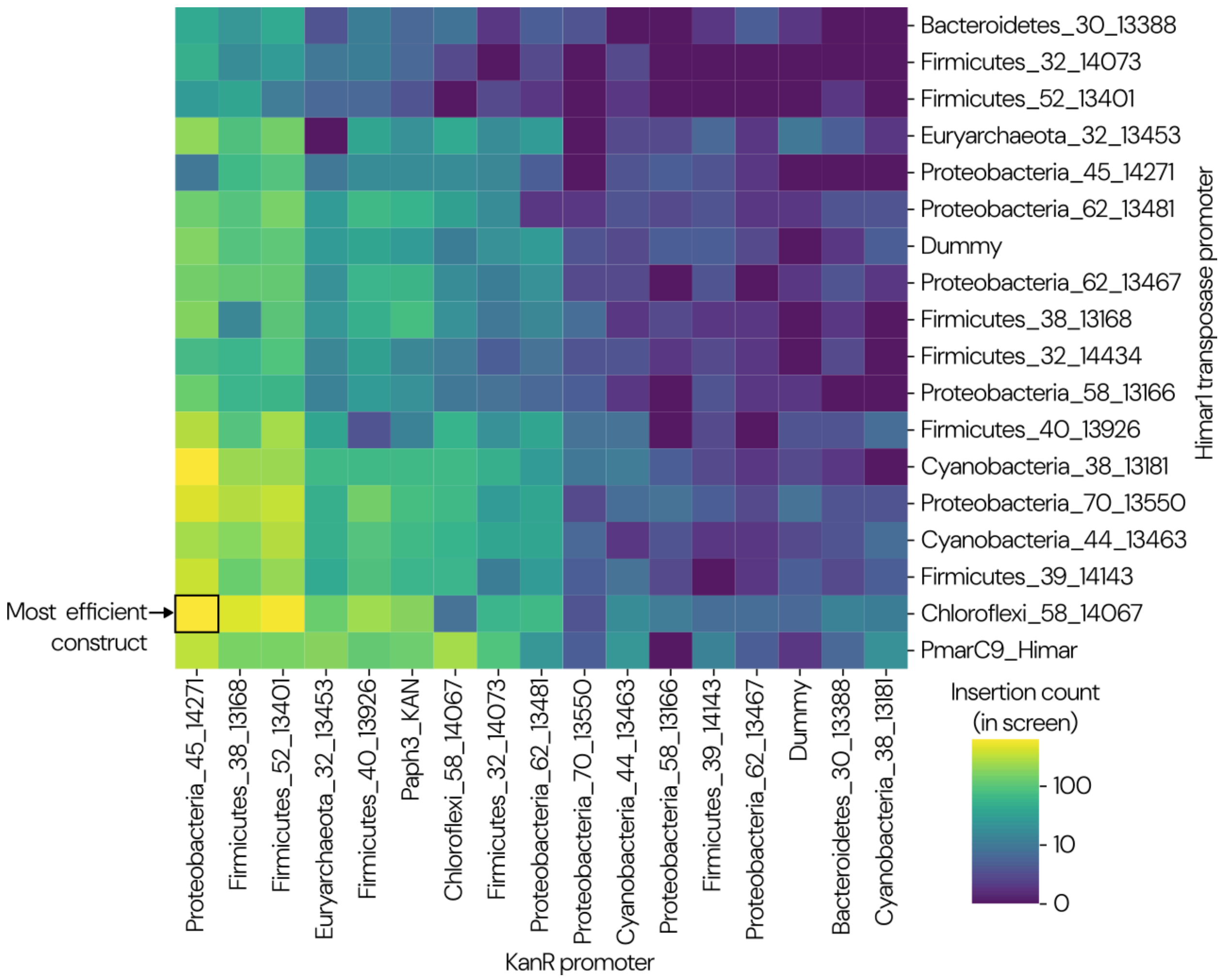


**Supplementary Figure 10. *Comamonas testosteroni* transposase and KAN selectable marker promoter efficiencies.**

Heat map plotting the total number of genomic insertion sites observed per unique combination of transposase and KAN gene promoter variants. The dataset shown was obtained from the automated 96-well plate format bacterial conjugation delivery of Himar1-only transposon variant libraries (**Figure 2A**). The combination of promoters that yielded the highest total number of insertions (labelled "Most efficient construct") was used to generate a genome-wide transposon mutant library in *C. testosteroni*.

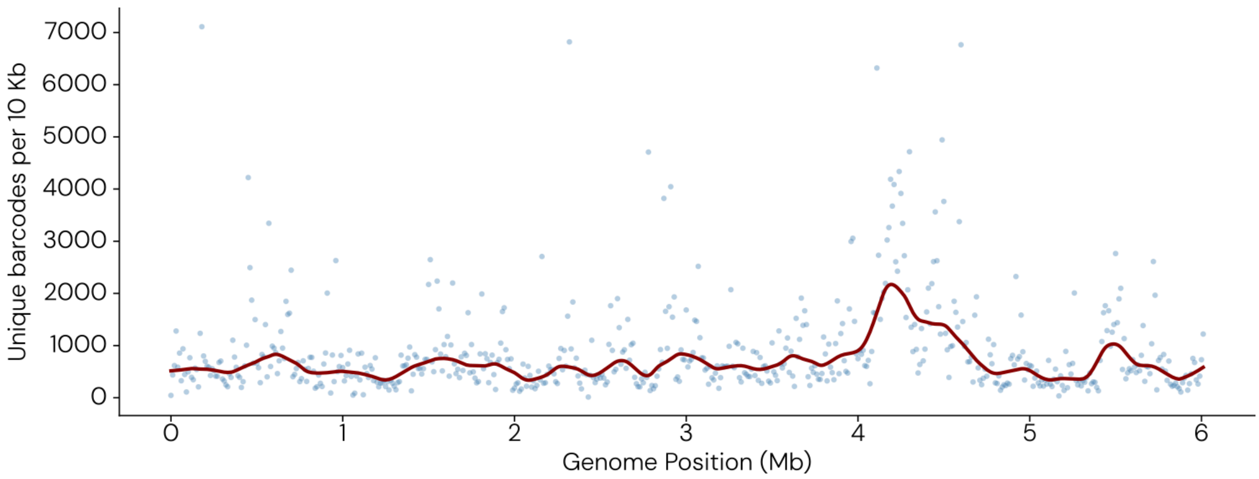


**Supplementary Figure 11. Genome-wide bias in insertion site frequency.**

The number of unique barcodes with identifiable insertion sites per 10 kb window plotted across the *C. testosteroni* genome.

#### Supplementary Data Files

**Supplementary Data Table 1. Promoter variants used in transposon variant libraries.**

**Supplementary Data Table 2. Details of transposon destination vector plasmids used in transposon variant libraries.**

**Supplementary Data Table 3. Details of the composition and construction of all transposon variant libraries described in this study.**

**Supplementary Data Table 4. Details of the composition and construction of individual transposon variant plasmids described in this study.**

**Supplementary Data Table 5. Strains, media, and antibiotics screened in this study.**

**Supplementary Data Table 6. Promoter efficiencies and insertion counts for all samples sequenced by Tn-seq.**

**Supplementary Data Table 7. Abundance, fitness, and significance values for all *C. testosteroni* genes during the terephalate growth experiment.**

**Supplementary Data Table 8. Abundance, fitness, and significance values for all C. testosteroni genes during the 4-hydroxybenzoate growth experiment.**
